# Evolutionary processes driving the rise and fall of *Staphylococcus aureus* ST239, a dominant hybrid pathogen

**DOI:** 10.1101/2021.01.10.426095

**Authors:** Jacqueline L. Gill, Jessica Hedge, Daniel J. Wilson, R. Craig MacLean

## Abstract

*Staphylococcus aureus* ST239 has been one of the most successful epidemic MRSA strains, and one of the leading causes of healthcare-associated MRSA infections. Here we investigate the evolution of ST239 using a combination of computational and experimental approaches. ST239 is thought to have emerged by a large scale chromosomal replacement event in which an ST8 clone acquired approximately 600 kb of DNA from an ST30 clone. Analysis of large-scale genomic data sets allowed us to confirm and refine the model of the origin of ST239. Importantly, we found that ST239 originated between the 1920s and 1945, implying that this MRSA lineage evolved at least 14 years before the clinical introduction of methicillin. Molecular evolution within ST239 has been dominated by purifying selection, although we found some evidence that the acquired region of the genome has evolved rapidly as a result of relaxed selective constraints. Crucially, we found that ST239 isolates have low competitive ability relative to both ST30 and ST8, demonstrating that this hybrid lineage is characterized by low fitness. We also found evidence of positive selection in a small number of genes involved in antibiotic resistance and virulence, suggesting that ST239 has evolved towards an increasingly pathogenic lifestyle. Collectively, these results support the view that low fitness has driven the recent decline of ST239, and highlight the challenge of using evolutionary approaches to understand the dynamics of pathogenic bacteria.

## Introduction

Antimicrobial resistance (AMR) in pathogenic bacteria has created a healthcare crisis by increasing the costs and mortality rates associated with bacterial infections. In many important pathogens, the rise of resistance has been driven by the epidemic spread of a small number of very successful clones, often those that have successfully acquired a range of resistance genes by horizontal gene transfer. One important challenge at the moment is to understand the evolutionary processes that give rise to these AMR superbugs^1^, such as *Staphylococcus aureus* sequence type ST22^2^, *Escherichia coli* ST131^3^ and *Klebsiella pneumonia* ST258^4^.

*S. aureus* is an important commensal pathogen that provides a clear illustration of the AMR crisis. Approximately 1/3 of humans show persistent asymptomatic carriage of *S. aureus*, mainly in the nares, but this bacterium is capable of causing serious invasive infections at a number of sites in the body^5^. Antibiotic use has driven the epidemic spread of waves of resistant *S. aureus* strains; in particular, epidemic strains of methicillin-resistant *S. aureus* (EMRSAs) are now a global problem in healthcare settings and a growing problem in the community. For example, ST239 (a member of clonal complex CC8) is a major EMRSA strain that has been the causative agent of multiple epidemics in healthcare settings around the globe. The first ST239 isolates were collected in the late 1970s in Australia^6^. This was followed by the first EMRSA strain, also an ST239 and known as EMRSA-1, which emerged in the UK in 1981^7^. Between 1987 – 1988, over 40% of MRSA isolates collected in England and Wales were ST239 EMRSA-1^8^, and ST239 became highly prevalent around the globe. By the mid-2000s, ST239 was the predominant MRSA strain in Asia, causing up to 90% of hospital-acquired MRSA within a region accounting for over 60% of the world’s population^9^. More recently, reports from around the globe show that levels of ST239 MRSA have been decreasing dramatically, but the causes of this decline remain unclear^10 11 12^.

Robinson and Enright proposed that ST239 was formed as a result of a large-scale chromosomal replacement event in which an ST8 clone acquired approximately 550 kb from an ST30 clone by recombination^13^. Interestingly, five of the six known other cases of large-scale chromosomal replacement in *S. aureus* overlap to some extent with the acquired region of ST239^13 14 15 16^, suggesting that this region may be a recombination hotspot^17^. Large-scale chromosomal replacements have also been detected in a number of important AMR pathogens outside of *S. aureus*, including *K. pneumonia* ST258, *Campylobacter coli* ST1150 and *Streptococcus agalactiae*^18 19 20^. In ST239, this acquired region also contains a large (60 kb) SCC*mec*-III element that confers resistance to a broad spectrum of antibiotics and heavy metals, and ST239 is thought to have spread in healthcare settings around the world as a result of the selective advantage of the resistance provided by this acquired element^21^.

Although subsequent studies have provided good support for the Robinson and Enright model, important questions regarding the origin of ST239 remain unresolved^22 23 24^. All of the 2,979 ST239 genomes in the Staphopia database that are associated with a typable SCC*mec* element carry a type III SCC*mec* element, suggesting that this element was acquired early in the evolutionary history of ST239. However, the SCC*mec*-III element has not been identified in any ST30 genomes, including all 1,896 ST30 genomes in the Staphopia database. The absence of SCC*mec*-III in ST30 suggests that the true ancestor of ST239 may be a closely related lineage that has not been considered in previous studies of ST239. Secondly, the origins of ST239 remain unclear. It is often assumed that antibiotic resistant pathogens evolve in response to treatment; in this case, it has been argued that the introduction of methicillin into clinical practice in the 1960s drove the evolution of ST239. However, recent work has shown that some other MRSA STs from CC8 pre-dated the clinical introduction of methicillin^25^, raising the possibility that ST239 may also have a deeper evolutionary history.

Finally, it is unclear why the prevalence of ST239 has recently declined. Acquiring DNA is usually associated with fitness costs^26 27 28^, and many studies have shown fitness costs associated with acquired resistance genes^29 30 31^, including large SCC*mec* elements^32 33^. Given these costs, it is possible that the recombination event(s) that gave rise to ST239 created a low fitness ‘hybrid’ pathogen, although this has not previously been investigated in detail.

In this paper, we use a combination of computational and experimental techniques to investigate the underlying evolutionary processes that have driven the rise of ST239. First, we assemble a diverse collection of ST239 genome sequences to reconstruct the evolutionary history of this lineage and to test the Robinson and Enright model. We then use competition assays to test the hypothesis that ST239 has low fitness, and to explore the link between AMR and fitness. Finally, we use population genetic approaches to understand how selection has operated in ST239, and to identify key genes involved in adaptive evolution in this lineage.

## Results and discussion

### Reconstructing the origin of ST239

The evolutionary drivers that contributed to the global emergence of the epidemic multi-drug-resistant MRSA strain ST239 are not well understood, however it has been suggested that the introduction of methicillin into clinics in the early 1960s may have been a contributing factor^34^. To reconstruct the evolutionary history of ST239, we assembled a collection of 96 ST239 genomes from isolates that were collected from diverse geographic locations and time points, deliberately avoiding over-representing isolates from intensively sampled ST239 outbreaks (e.g.^35^ and ^23^) (Supplementary Table 1A).

We constructed a pan-genome of 3,337 ST239 genes, of which 1,889 were present in 99 – 100% of all strains, resulting in a core genome length of 1,754,805 bp, representing a greater number of ST239 isolates from varied locations and dates than in previous evolutionary studies of ST239^36 37 38^. After identifying and excluding sites involved in recombination^39^, 3,696 core variant sites remained, from which we reconstructed a phylogeny by maximum likelihood (Figure 1A). The phylogeny had strong bootstrap support, and the long branch lengths were suggestive of a diverse set of ST239 isolates^40 34^. The clustering of isolates into geographically distinct clades is consistent with the population structure observed by Harris (2010), and by Castillo-Ramirez (2012), who identified strong geographical clustering of ST239 sequences on continental, national and city scales^37 34^. In this study, isolates from Oceania, Asia and South America formed distinct clades, with rare exceptions. In contrast, North American and European isolates were dispersed throughout the tree.

**Figure 1A.**
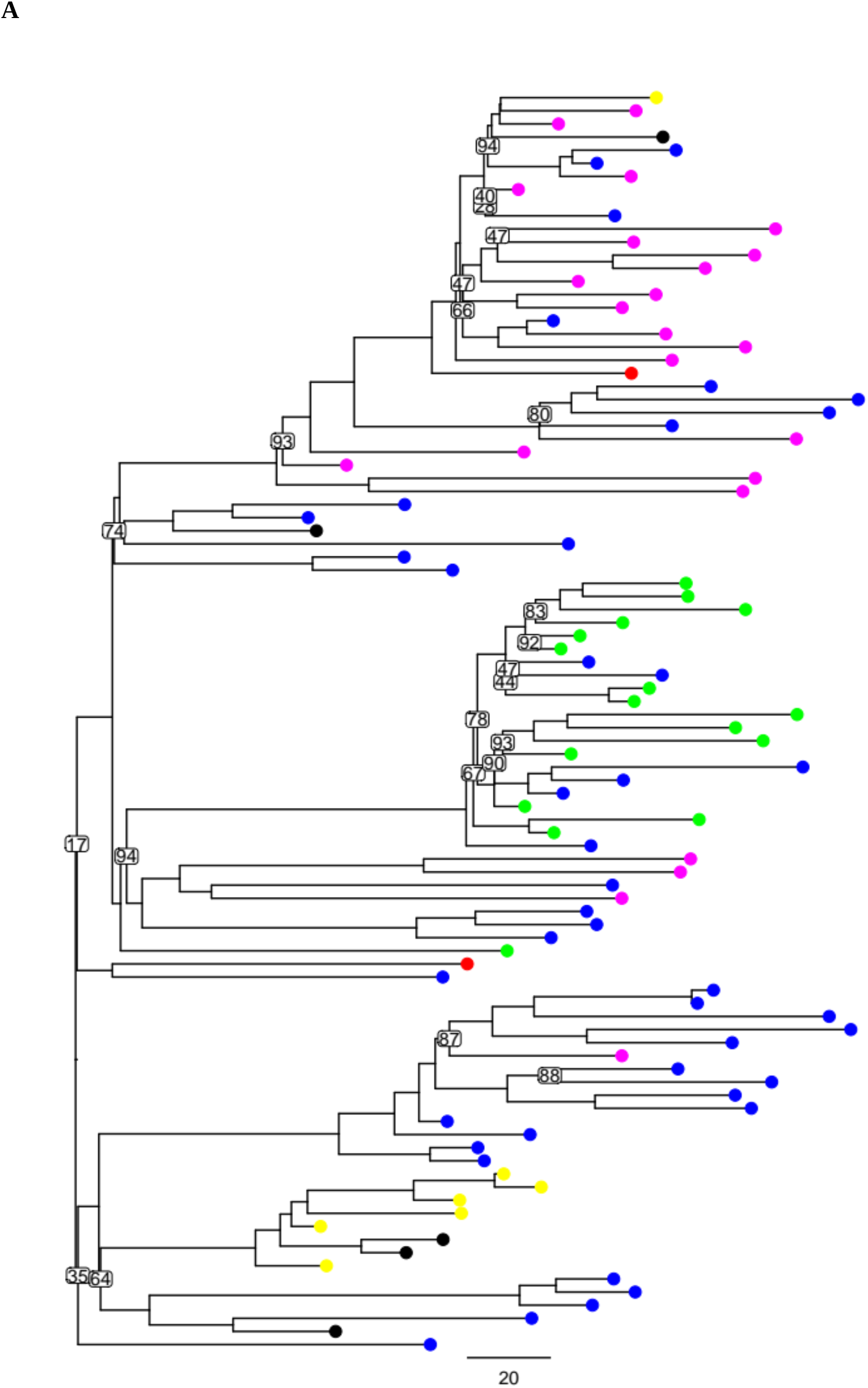
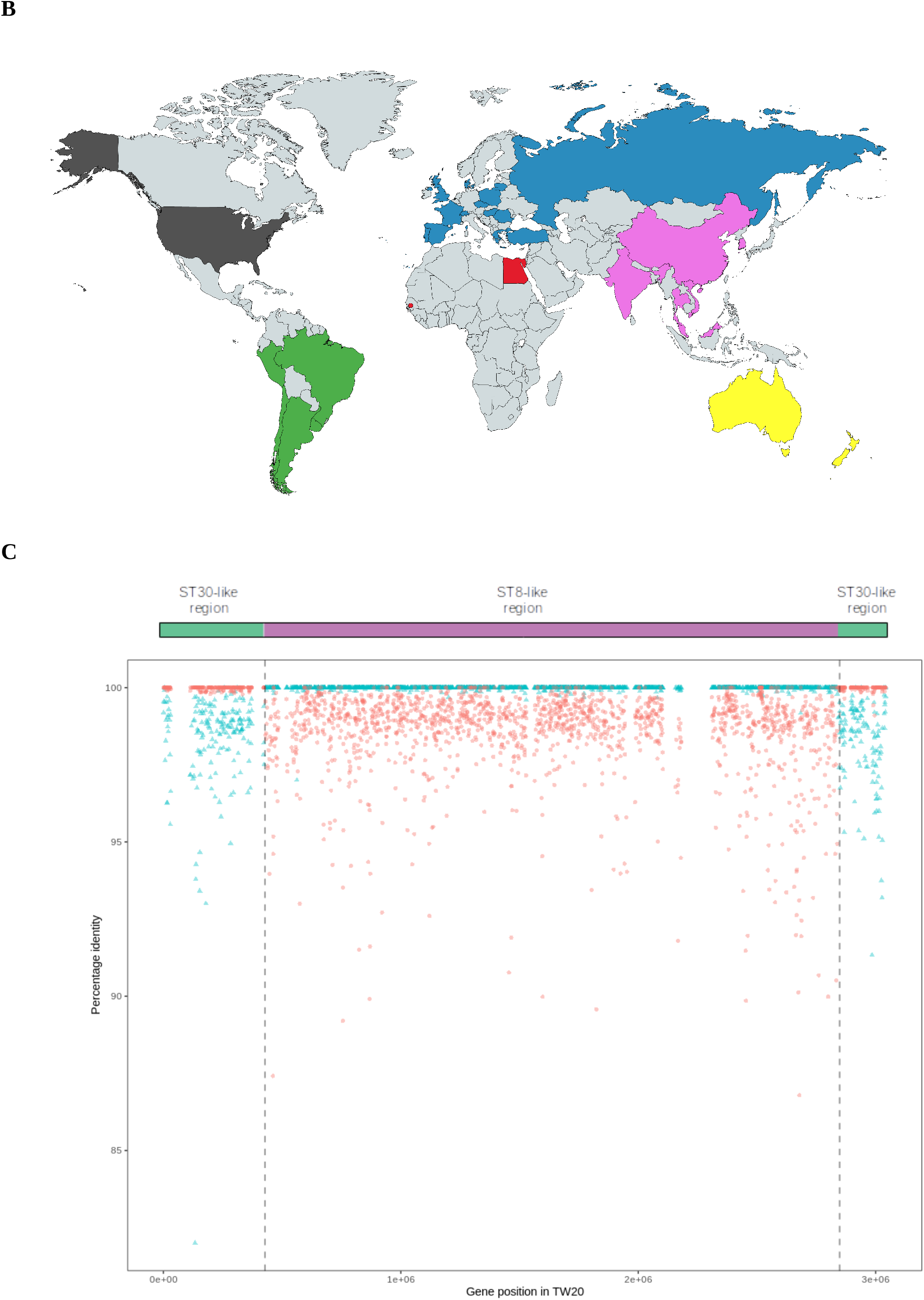
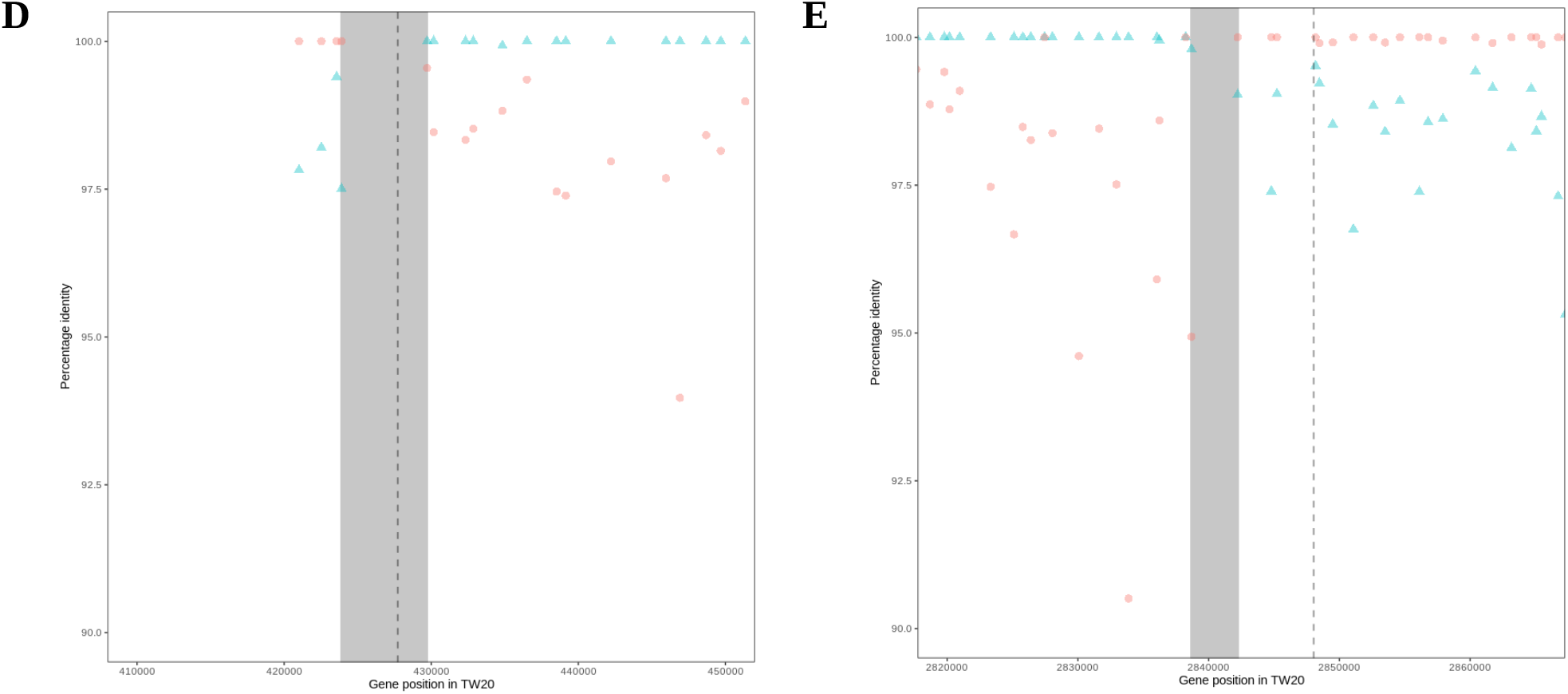
Out-group rooted maximum-likelihood phylogeny of the 96 ST239 genomes. Distances are shown in in SNPs/Mb and bootstrap support values below 95 are shown for branches. Isolates are colour coded according to geographical origin, as displayed on panel B (Blue, Europe; Pink, Asia; Yellow, Oceania; Green, South America; Black, North America; Red, Africa). **1C.** BLAST DNA percentage identity of the ST8 (blue triangle) and ST30 (red circle) consensus core gene sequences, compared to the ST239 consensus core gene sequences (consensus sequences were formed from 96 ST239 genomes, 111 ST8 genomes and 57 ST30 genomes, for 1,980 core genes that were shared between all 264 genomes). In the ST239 genome, the acquired region spans the origin of replication, and hence is split between the beginning and the end of the linearized genome. The hybrid boundaries estimated by Castillo-Ramirez *et al*. (2011) are highlighted with vertical dashed grey lines. **1D.** Close up of the first boundary of the large chromosomal replacement event. **1E.** Close up of the second boundary of the large chromosomal replacement event. The hybridisation boundaries estimated by Castillo-Ramirez *et al*. (2011) are highlighted with vertical dashed grey lines, and the hybridisation boundary ranges used in this study are highlighted by a grey rectangle. Each data-point represents a single gene. The gene positions correspond to the genomic position within the ST239 reference genome.

We estimated a time to the most recent common ancestor (MRCA) of these ST239 sequences by fitting four evolutionary models using BEAST (Supplementary Table 2A)^41^. The time to MRCA was consistent between all four models, with no significant difference between the models identified through Bayes Factor analysis. Therefore, the time to MRCA from the simplest model (strict molecular clock, constant population size) is recorded here, as 1940.1 (95% Highest Posterior Density (HPD) intervals: 1934.7 – 1945.5). Similar results were obtained using BactDating (Supplementary Table 2B). Although there is some uncertainty in these estimates, these models predict that the origin of ST239 predated the clinical introduction of methicillin in 1959^42^ by more than 10 years.

### Identifying the donor of the acquired region of the ST239 genome

Robinson and Enright initially proposed that ST30 was the closest known ancestor of the ST239 acquired-region^13^, and they were able to identify potential boundaries of the recombination event that gave rise to ST239^13^. However, their conclusions were based on the partial sequencing of a small number of genes from representative isolates. To test the Robinson-Enright model using a large collection of genomes, we extracted the core genes shared by >99% of the isolates from the ST239 collection (*N* = 96), combined with additional collections of diverse ST8 (*N* = 111) and ST30 (*N* = 57) genomes. A total of 1,980 core genes were identified that were shared between >99% of all 264 genomes. Within each ST (ST239, ST8 and ST30), we generated a consensus sequence for each of these genes, and calculated the percentage similarity between each ST239 gene and its homolog in either ST30 or ST8 (Figure 1C).

There was a clear distinction between regions of the ST239 genome that were closely related to ST8 and ST30, allowing us to clearly differentiate between the backbone and acquired regions of the ST239 genome (Figure 1D, E). This was consistent with Holden *et al*., who compared an ST239 genome sequence with a CC30 complete genome sequence, and found that the acquired region was more closely related to CC30 by a shift of roughly 1% in DNA percentage identity compared to the backbone region^21^.

Although the acquired region of the ST239 genome is similar to ST30, it is possible that the true ancestor of this region was a closely related lineage of *S. aureus* that was not considered in previous analyses of ST239. For example, all of the 2,979 ST239 isolates in the Staphopia database that are associated with a typable SCC*mec* element carry a type III SCC*mec* element, but this element has not been identified in any of the 1,896 ST30 genomes in the Staphopia database^43^, suggesting that ST30 may be a sister group, rather than the true ancestor of the acquired region (Supplementary Table 3). To systematically search for the ancestor of the acquired region, we used BIGSI to screen 447,833 bacterial and viral raw-read and assembled genomes for short sequences that match those found in the acquired region of the ST239 genome^44^. Sequences from isolates that were closely related to the acquired region of the ST239 genome were mapped to the acquired region of the ST239 reference genome (NCBI accession number FN433596), and assembled into a phylogeny alongside the acquired region of the 96 ST239 genomes (Figure 2A). Crucially, we found that all ST239 isolates share a recent common ancestor with five ST30 isolates dating from the 1950s and 1960s (and one ST30 isolate of unknown origin), and this branch is well-supported by bootstrapping. These five ST30 isolates (Supplementary Table 4) were all from the penicillin-resistant *S. aureus* phage type 80/81 clone, which caused serious healthcare-associated and community-associated infections worldwide, and was largely eliminated in the 1960s as a result of the introduction of methicillin^42 45 46^.

**Figure 2A.**
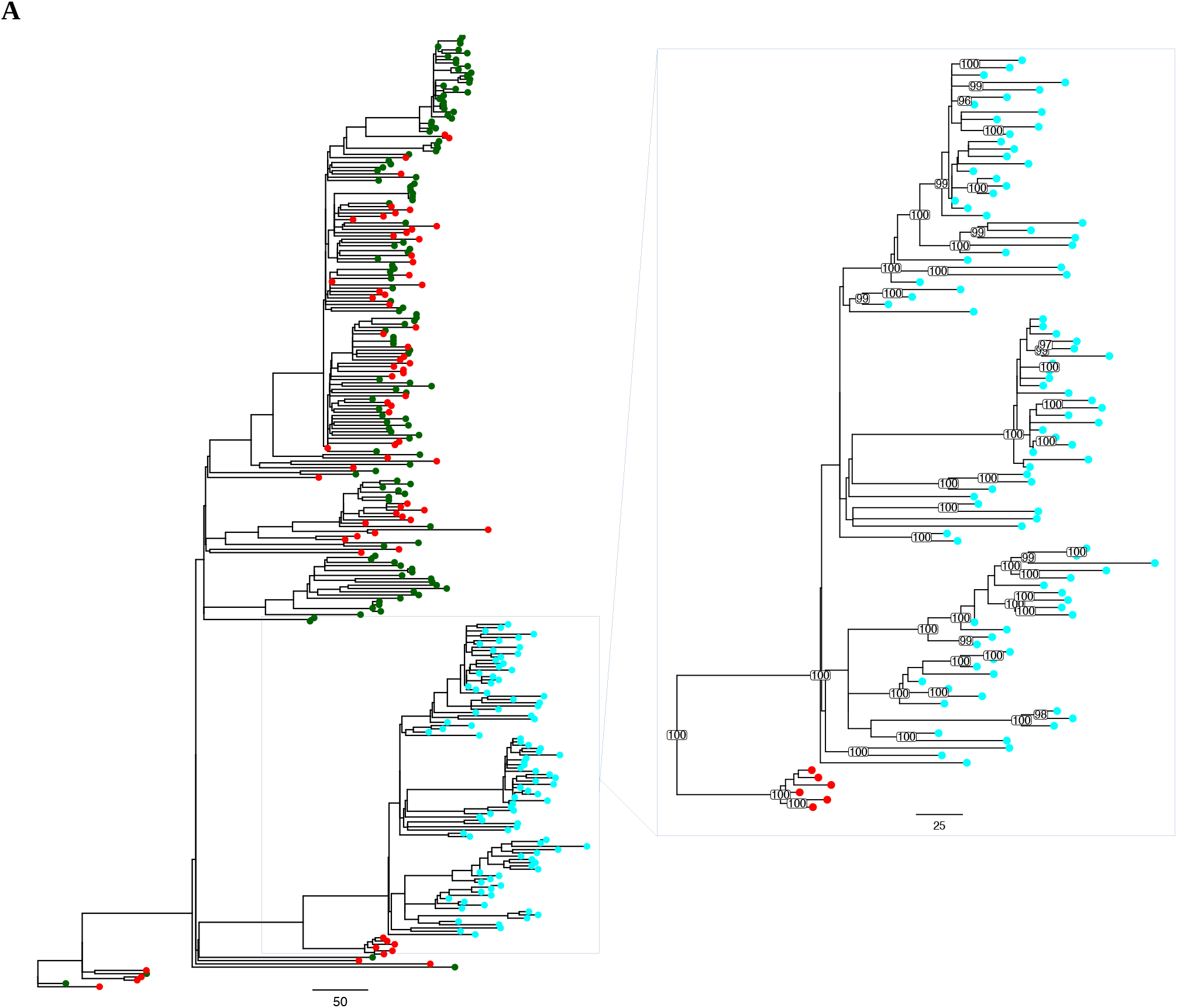
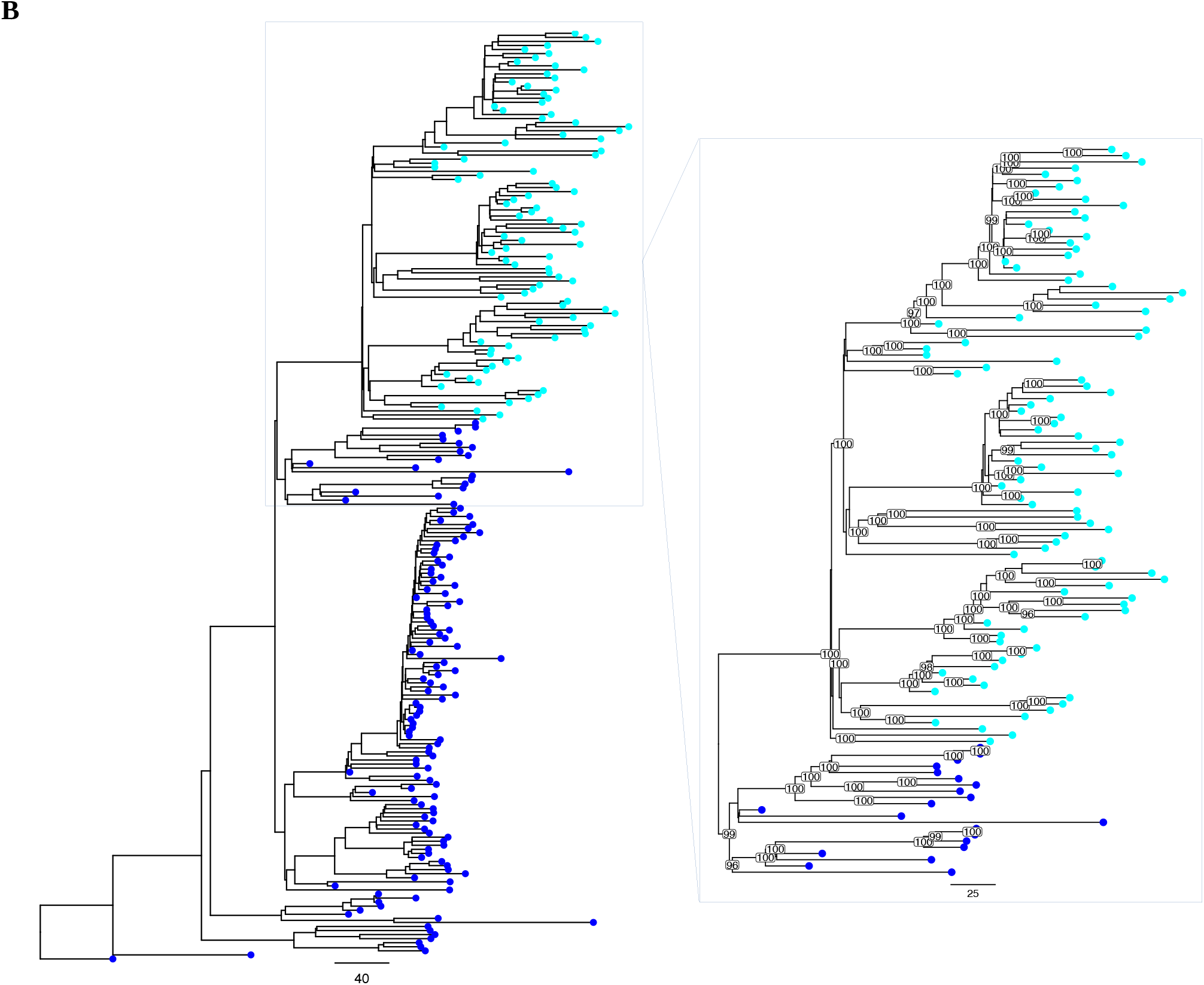
Outgroup-rooted maximum likelihood phylogeny of the acquired region of the ST239 genomes, and the most closely related non-ST239-like isolates from BIGSI analysis. Distances are shown in in SNPs/Mb. ST239 isolates are in cyan, ST30 isolates in red and other STs are in green. The zoom panel contains the ST239 clade and the most closely related non-ST239 isolates, with bootstrap support values above 95 shown for branches. **2B.** Outgroup-rooted maximum likelihood phylogeny of the backbone region of the ST239 genomes, and the most closely related non-ST239-like isolates from BIGSI analysis. Distances are shown in in SNPs/Mb. ST239 isolates are in cyan and ST8 isolates are in blue. The zoom panel contains the ST239 clade and the most closely related non-ST239 isolates, with bootstrap support values above 95 shown for branches.

Interestingly, none of the phage type 80/81 genomes that are closely related to ST239 contain an SCC*mec* element. The ubiquity of SCC*mec*-III in ST239 genomes suggests that the SCC*mec*-III element was acquired by the ancestor of the acquired region of ST239 following the divergence of this lineage from the phage type 80/81 lineage. However, it is possible that the SCC*mec*-III element was secondarily acquired following the chromosomal replacement that gave rise to ST239. To test the secondary acquisition hypothesis, we estimated the time to MRCA of the SCC*mec*-III elements in the collection of ST239 isolates (Supplementary Figure 1, Supplementary Table 2A). However, there was considerable uncertainty in this estimate due to the small size of the SCC*mec*-III element (60 kb) and the high rate of recombination in this region of the genome. Given, these limitations, this analysis had limited power to reject the null hypothesis that the SCC*mec*-III element was acquired as part of the initial chromosomal replacement event that gave rise to ST239.

As a final test of the Robinson-Enright model, we used BIGSI to identify the closest known ancestor of the backbone region of the ST239 genome. Consistent with the Robinson and Enright model, this analysis identified ST8 as the closest ancestor of the ST239 backbone region^13^. Specifically, the ancestor of ST239 was part of a diverse lineage of ST8 that has been isolated across multiple continents over the last 70 years (Figure 2B, Supplementary Table 4).

Dating the MRCA of ST239 isolates places an upper bound on the origin of ST239, but it is possible that ST239 originated prior to the MRCA of contemporary isolates. To place a lower bound on the origin of ST239, we estimated times to the MRCA of the acquired and backbone regions of ST239 with their respective ST8 and ST30 ancestors. The MRCA of the acquired region and ST30 was estimated as 1900.3 – 1926.5 (95% HPD) and MRCA of the backbone region and ST8 was estimated as 1924.5 – 1934.7 (95% HPD). The overlap of these two estimates is encouraging, and this analysis suggests that ST239 is unlikely to have originated prior to the 1920s.

### Recombination

Given that recombination created the ST239 lineage, it is possible that recombination has also played an important role in the subsequent evolution of ST239. To investigate this idea, we used ClonalFrameML to identify regions of recombination within the collection of 96 ST239 genomes (Figure 3, Supplementary Figure 2). Ten isolates from the TW20-like clade (Supplementary Table 5) were excluded from this analysis due to high sequence similarity to the reference sequence that all ST239s were mapped to.

**Figure 3.**
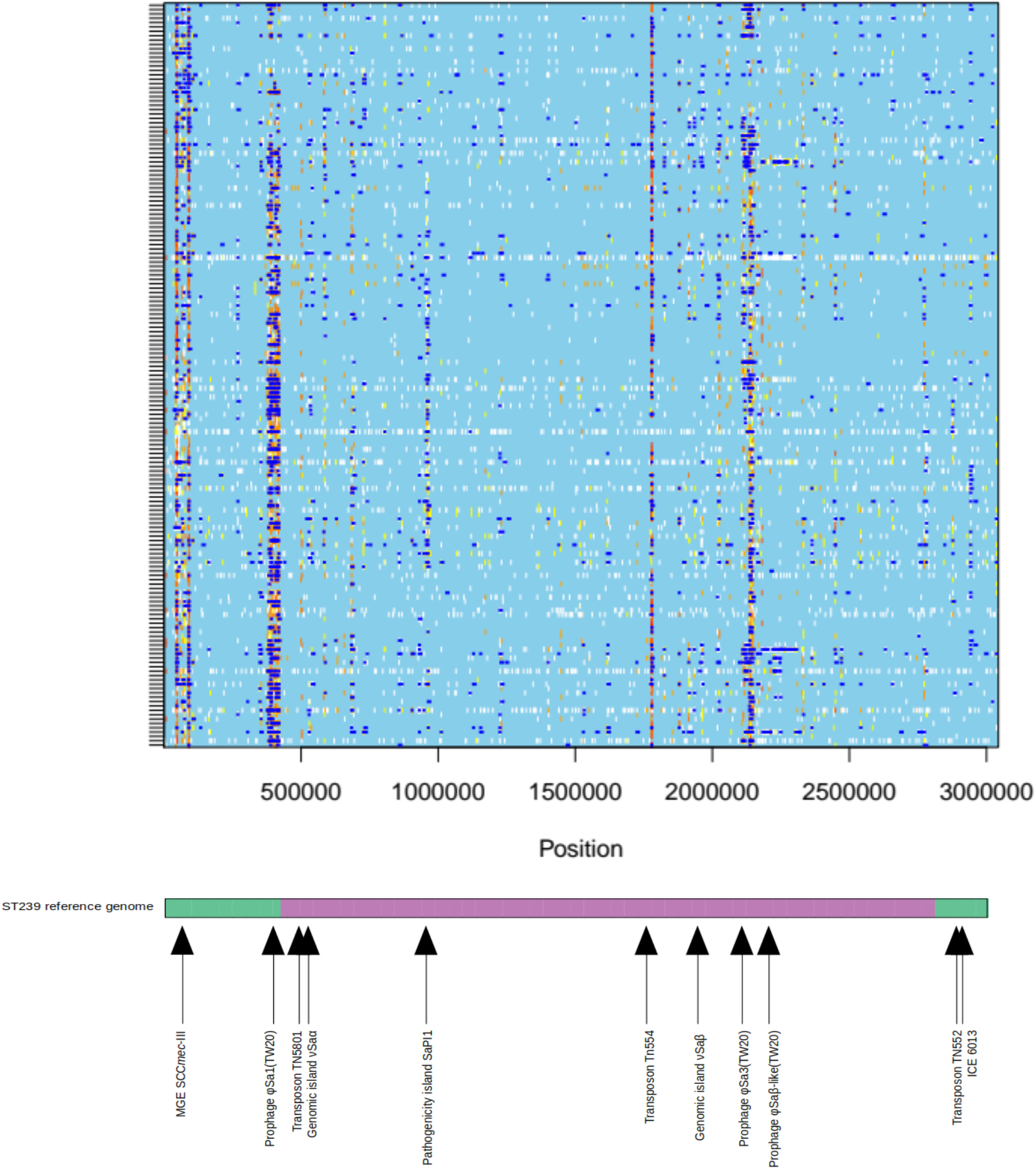
Recombination plot of the ST239 genomes compared with the ST239 TW20 reference genome. Dark blue regions represent potential sites of recombination. Sites that are non-polymorphic are shown in light blue. Polymorphic sites are shown in a colour indicating their level of homoplasy; white is no homoplasy and the range from yellow to red is increasing degrees of homoplasy. The positions of known MGEs and the two inherited regions estimated by Holden *et al*. and Castillo-Ramirez *et al*. are highlighted along the ST239 reference genome beneath the plot; green, acquired region; magenta, backbone region.

The ratio of rates of recombination and mutation (*R*/*θ*) was 0.41, and the ratio of effects of recombination and mutation (*r/m*) was 2.37. Therefore, recombination events occurred 2.5 times less often than mutations, however, because each recombination event introduced an average of 6.0 substitutions, recombination overall was responsible for 2.4 times more substitutions than mutations^47^. In line with previous analyses, we found evidence of recombination hotspots in regions of the genome containing mobile genetic elements^24^. Recombination was particularly frequent in the region surrounding the φSA1 prophage, suggesting that this element, which borders the acquired region, may have played a key role in the recombination event that gave rise to ST239. Notably, the SCC*mec*-III element is also a hotspot for recombination.

### Evolutionary consequences of large-scale chromosomal replacement

To begin to understand the evolutionary consequences of large-scale chromosomal replacement, we calculated the evolutionary rate of the backbone and acquired regions of the ST239 genome using BEAST (Supplementary Table 2A)^38^. After removing recombination, the overall genomic substitution rate was estimated as 1.205 x 10^−6^ SNPs/site/year (95% HPD 1.13 × 10^−6^ – 1.28 × 10^−6^ SNPs/site/year), which is similar to the substitution rates of other EMRSAs^2 48 49 50^. Interestingly, the substitution rate of the acquired region, 1.515 × 10^−6^ (1.32 × 10^−6^ – 1.71 × 10^−6^ SNPs/site/year; 95% HPD), was 2% – 51% more rapid than the backbone, 1.21 (1.13 × 10^−6^ – 1.29 × 10^−6^ SNPs/site/year; 95% HPD). On the one hand, the rapid evolution of the acquired region could be a signature of rapid adaptive evolution, perhaps to overcome the costs associated with horizontal gene transfer. Alternatively, it is possible that the acquired region has evolved at a high rate due to weak selective constraints in this region of the genome.

To understand how selection has acted in the two genetic regions of the ST239 genome, we used the McDonald-Kreitman test to compare patterns of polymorphism and divergence in the backbone and acquired regions of the genome^51^. The McDonald-Kreitman net neutrality index (*N*) indicates whether selection is overall purifying (*N* >1) or positive (*N* <1)^52^.

In this analysis, we compared the acquired region of the 96 ST239 genomes with the homologous region in the previously defined collection of 57 ST30 genomes. Additionally, we compared the backbone region of the 96 ST239 genomes with the homologous region in the previously defined collection of 111 ST8 genomes. Only “core” genes that were shared by more than 99% of all ST239, ST8 and ST30 genomes in the collections were included. The divergence between ST239 and ST30 in the acquired region (0.326 substitutions/Mb) was much greater than the divergence between ST239 and ST8 in the backbone region (0.188 substitutions/Mb), reflecting the fact that ST239 is more closely related to ST8 than ST30 in the respective regions. Divergence between STs at non-synonymous sites was low relative to levels of within-ST polymorphism, demonstrating an overall trend towards purifying selection in ST239 (Table 1; *N* >1). However, we did not find any difference in the net neutrality index between the backbone (*N* = 2.06; 95% C.I. *N* = 0.92 – 2.51) and the acquired region (*N* = 1.52; 95% C.I. *N* = 1.34 – 3.19).

**Table 1.**
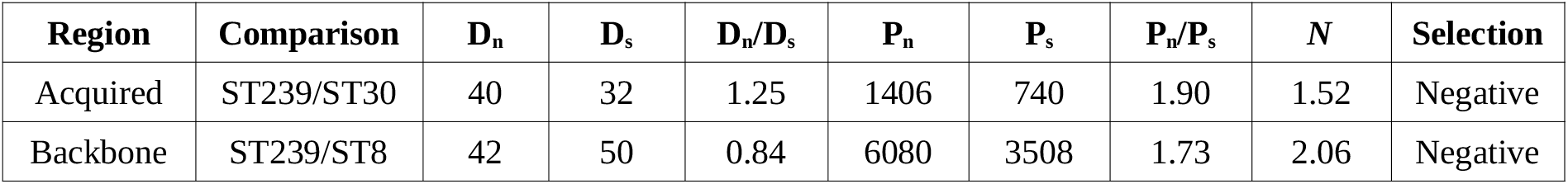
McDonald-Kreitman net neutrality index for the two genetic regions of ST239. Fixed non-synonymous substitutions (Dn); fixed synonymous substitutions (Ds); non-synonymous polymorphisms (Pn); synonymous polymorphisms (Ps); McDonald-Kreitman net neutrality index (N).

One weakness of this approach in this context is that the backbone and acquired regions of the ST239 genome have to be compared to different outgroups. Given that signatures of purifying selection should become stronger over time, it could be argued that the McDonald-Kreitman test is biased towards detecting purifying selection in the acquired region of the genome, which was compared to a more divergent outgroup. To further test the idea that selection on the acquired region has been weak, we calculated the ratio of GC → AT to AT → GC substitutions in each region of the ST239 genome (Table 2). Spontaneous mutation in *S. aureus* is biased towards GC → AT transitions, and regions of the genome that are subject to weak selective constraints are therefore expected to have a high ratio of GC → AT/AT → GC substitutions^53^. In this case, the GC → AT/AT → GC ratio in the acquired region was significantly greater than that of the backbone region (Fisher’s exact test, odds ratio = 1.15, *P* = 0.0303). This analysis, which uses data on substitutions that have occurred during the diversification of ST239, provides evidence of relaxed selective constraints on the acquired region of the ST239 genome.

**Table 2.**
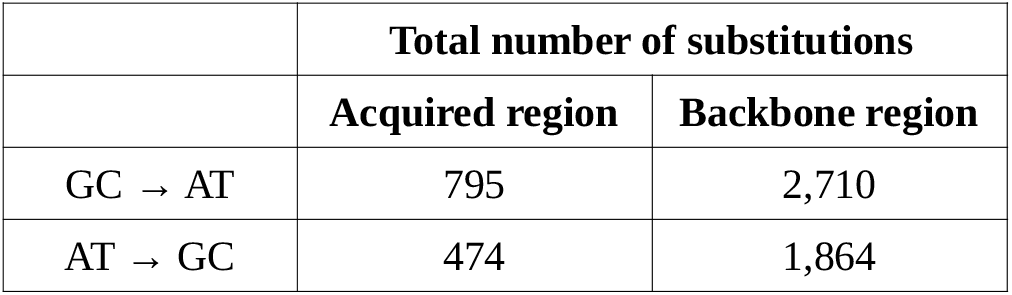
Total number of GC → AT and AT → GC substitutions in the acquired and backbone regions of the ST239 genome.

### Fitness costs of chromosomal replacement

To understand the fitness effects of chromosomal replacement more directly, we measured the competitive ability of a collection of *S. aureus* isolates *in vitro* (Supplementary Figure 3). Our collection of isolates included divergent isolates of ST239 (*N* = 4), ST8 (*N* = 6) and ST30 (*N* = 5) that were deliberately chosen to avoid including clonal isolates within STs. If chromosomal replacement is costly, then we would expect ST239 isolates to have low competitive ability relative to ST8. The additional ST30 isolates provided a useful reference point to compare fitness values. Isolates were directly competed against each other in three different culture media; Tryptic Soy Broth (TSB), Brain-Heart Infusion (BHI) broth, and Porcine Serum (PS). These media impose different stresses that mimic some of the challenges encountered by *S. aureus* in clinical environments.

We deep sequenced (142 – 211x depth) each competition mixture before and after growth, and estimated changes in the relative abundance of each isolate by quantifying the relative abundance of isolate-specific SNPs during competition. Competitive ability varied between isolates, and we found a strong statistical interaction between ST and media, which reflects the fact that the average competitive ability of the three STs varied across media (Figure 4 A – C; Table 3; Supplementary Table 6A). For example, we did not find any evidence of low fitness associated with ST239 in porcine serum. In spite of this variation in fitness, we found a significant difference in competitive ability between STs. Crucially, we found that ST239 had lower competitive ability than both ST8 and ST30 (Figure 4D; post-hoc Tukey test *P* <0.05) The low fitness of ST239 relative to the ST8 is consistent with the idea that chromosomal replacement carries a long-term fitness cost. This hypothesis is further supported by the high fitness of ST30, which suggests that the difference in fitness between ST239 and ST8 reflects low fitness of ST239, rather than high fitness of ST8.

**Figure 4.**
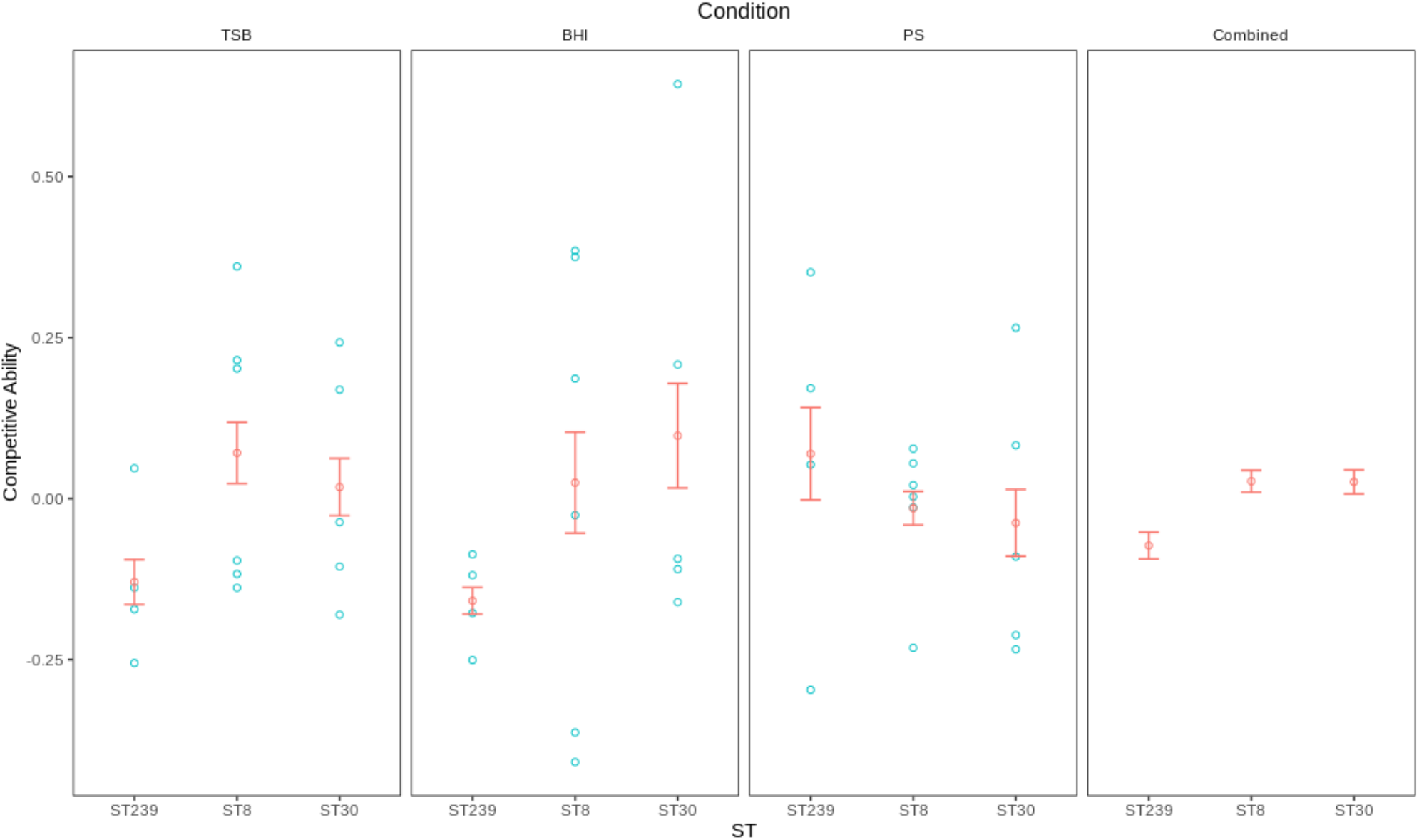
Competitive ability. Plotted points show the competitive ability of *S. aureus* isolates (blue circles) and the mean competitive ability of each ST (red circles; +/− s.e.m; *N* = 4−6). The competitive ability of each isolate was measured in triplicate, and error for individual isolates was small (s.e = 0.045-0.001). ST239 isolates have reduced competitive ability relative to ST8 and ST30 in BHI and TSB, but not PS, as judged by a post-hoc Tukey test (*P* <0.05). The final panel shows the overall effect of ST on competitive ability (+/− s.e.) across all three media.

**Table 3.** Reduced ANOVA table of significant competitive ability effects.

| Effect | df | Sum of squares | Mean squares | F-value | *p*-value |
| --- | --- | --- | --- | --- | --- |
| ST | 2 | 0.260 | 0.130 | 8.33 | 0.0004 |
| Media x ST | 4 | 0.565 | 0.141 | 9.05 | <0.0001 |
| Isolate[ST] | 12 | 4.24 | 0.354 | 22.6 | <0.0001 |
| Error | 116 | 1.81 | 0.0156 | - | - |

As a complementary approach to measure fitness, we also measured the growth rate of the individual ST239, ST8 and ST30 isolates in TSB, BHI and PS (Supplementary Figure 4). There was significant variation in growth rate between isolates, media, and ST showing that this trait is very plastic (Supplementary Table 6B). Crucially, we found that ST239 isolates had significantly lower growth rate than ST8 isolates, providing further evidence of costs associated with chromosomal replacement.

### Parallel evolution

A recurring theme of studies of microbial evolution is that genes that are under strong positive selection evolve in parallel^54 55 56^. To better understand the selective pressures that have shaped the evolution of ST239, we compared the distribution of observed mutations per gene with a neutral model derived from the Poisson distribution, in which mutations are randomly distributed across genes. This analysis was carried out independently for the acquired and backbone regions of the ST239 genome to take into account the differences in substitution rate between these genomic regions. Only the 1,980 “core” genes that were shared between the previously defined ST239, ST30 and ST8 genome collections were included.

The number of mutations per gene differed from the Poisson expectation in both the acquired region (X^2^ = 19.158, *P* = 0.0014) and backbone region (X^2^ = 177.8, *P <*0.0001), showing that substitutions are non-randomly distributed across the ST239 genome. A subset of genes that show more evidence of parallel evolution than expected due to chance alone were defined as those that had 9 or more substitutions per gene. Our justification for this cut-off is that the Poisson distribution predicts that 1 or 2 genes in each region of the genome should have acquired 9 mutations or more due to chance alone, given an average of 2.53 substitutions per gene in the backbone region and 2.97 substitutions per gene in the acquired region. The proportion of genes showing evidence of parallel evolution did not differ between the backbone (20/1659 = 1.21% genes) and acquired regions (6/316 = 1.90% genes), indicating that genes under positive selection are evenly distributed across the ST239 genome (Chi-squared test, X^2^ = 0.9818, *P* = 0.3218). However, of the genes showing evidence of parallel evolution, those in the acquired region had a much larger proportion of substitutions (*N* = 154 substitutions, 15.75%) than the backbone region (*N* = 284 substitutions; 6.61%). The acquired region includes the *spa* gene (*N* = 62 substitutions) which is known to undergo strong selection mediated by the immune system. Even after excluding this gene, the remaining genes showing evidence of parallel evolution in the acquired region are significantly enriched for substitutions compared to those in the backbone (Chi-squared test, X^2^ = 13.111, *P* = 0.000294). The high rate of substitutions in these genes suggests that the acquired region has been a hotspot for adaptive evolution in the ST239 genome. Note that this analysis is robust to the overall elevated substitution rate of the acquired region because it is based on a sub-set of genes that show a high rate of substitution compared to other genes in the region.

Interestingly, many (*N* = 7) of the genes that show evidence of parallel evolution (Supplementary Table 7) are involved in resistance to antibiotics, including vancomycin/daptomycin (*walK*), fluoroquinolones (*grlA*), β-lactams (*ponA* and *mprF*) and rifampicin (*rpoB*). To test for elevated resistance at phenotypic level, we measured the resistance of our isolates to a broad panel of antibiotics that have activity against *S. aureus*. ST239 isolates were resistant to a greater number of antibiotics (Figure 5; Supplementary Table 8; mean = 7.25; s.e. = 0.75) than either ST30 or ST8 (One-Way ANOVA followed by Hsu’s test F_2,12_ = 5.63; *P* = 0.0187), highlighting the high levels of AMR associated with this lineage.

**Figure 5A.**
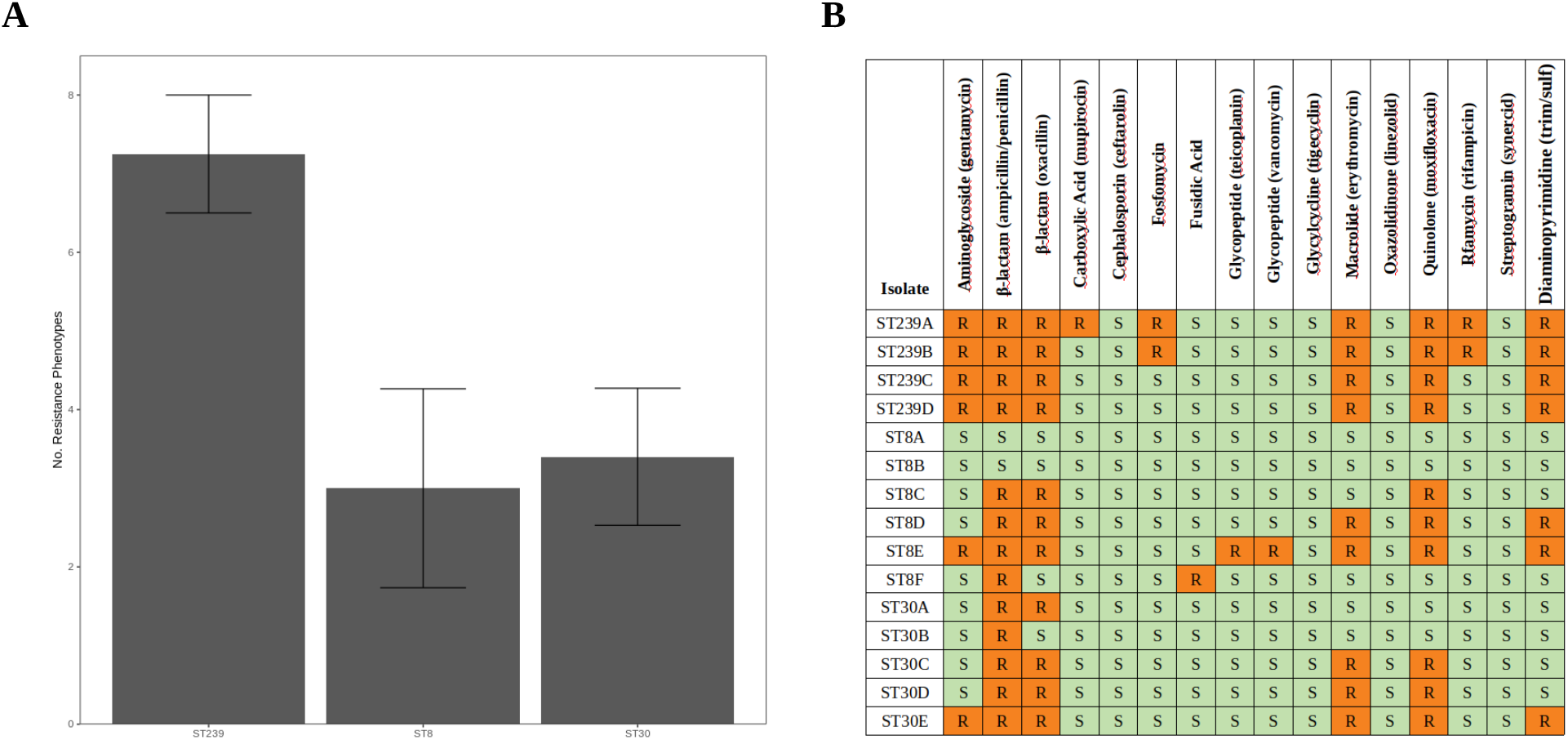
Mean (+/− s.e.m.; *N* = 4 – 6) number of AMR phenotypes for ST239, ST8 and ST30 isolates. **5B**. Heat map of AMR phenotypes for ST239, ST8 and ST30 isolates. Orange (R), resistant; green (S), susceptible.

## Conclusion

In line with previous work, our results support the hypothesis that ST239 was produced by a large scale chromosomal replacement event in which an ST8 clone acquired >600 kb of DNA from an ST30 clone^13 21^. We were able to refine this model by showing that the ancestor of the acquired region of the ST239 genome was most closely related to phage type 80/81 clones that were associated with the epidemic spread of penicillin resistance in the 1950s and 1960s^57^. The most parsimonious explanation for the presence of the SCC*mec*-III in ST239 is that this element was acquired by the ST30 ancestor of ST239, following the divergence of this lineage from phage type 80/81. However, our analysis had limited power to detect secondary acquisition of SCC*mec*-III. We estimate that ST239 originated between the 1920s and 1945, providing further evidence that MRSA pre-dated the clinical introduction of methicillin in 1959^25^. SCC*mec*-III provides resistance to first generation antibiotics that were used prior to the introduction of methicillin, such as tetracycline and erythromycin, and heavy metals, such as cadmium and mercury, that are used in disinfectants and biocides in healthcare settings^58 59^, suggesting that these resistance phenotypes may have provided ST239 with a selective advantage prior to the introduction of methicillin.

Fitness costs of laboratory-evolved antibiotic resistance have been demonstrated in many studies^28 60 61^, but estimates of the cost of resistance in pathogen populations have received less attention^62 63 64^. We found extensive variation in competitive fitness between *S. aureus* isolates and culture conditions. In spite of this variation, we found a clear overall trend towards low fitness in ST239 relative to ST8, providing good evidence of a fitness cost associated with the evolution of elevated antibiotic resistance. This hypothesis is further supported by epidemiological evidence. ST8 is primarily found in the community, where antimicrobial use is low, whereas ST239 has mainly been restricted to healthcare settings where antimicrobial use is high, suggesting that the low fitness of this ST has restricted the spread of ST239 into the community^65 21 66^. The acquisition of large SCC*mec* elements tends to generate a fitness cost, suggesting that the SCC*mec*-III element (which is the largest known SCC*mec* element) contributes to the low fitness of ST239 (see also^57^). Although we found some evidence that the acquired region of the ST239 genome is subject to relaxed selective constraints, the evolution of this region is dominated by purifying selection, suggesting that the chromosomal replacement may have had costs beyond those associated with the acquisition of SCC*mec*-III.

Experimental studies have found evidence that bacteria can adapt to the cost of gene acquisition through the process of compensatory evolution^25 27^. Although there are clear examples of compensatory adaptation in pathogenic bacteria (i.e.^67 68 69^), the prevalence of the compensatory evolution is unclear^59^. Although the evolution of ST239 has been dominated by purifying selection, we found evidence of positive selection in genes that are implicated in antibiotic resistance, virulence and metabolism. Notably, many of the genes that show clear hallmarks of lineage-specific positive selection in the ST239 genome are associated with resistance to antibiotics that have been used to treat MRSA infections, such as ciprofloxacin (*grlA*), vancomycin (*walK*), and rifampicin (*rpoB*). These patterns of parallel evolution suggest that the ST239 has evolved to increase antibiotic resistance and virulence, rather than to overcome the costs associated with chromosomal replacement and SCC*mec*-III acquisition.

Bacterial recombination is typically associated with the exchange of short DNA sequences between closely related strains or species^70^, and the large-scale chromosomal replacements that have been detected in pathogenic bacteria are conspicuous exceptions to this overall trend^18 19 20^. Our results support the idea that ST239 is a ‘hopeful monster’ that has declined in prevalence due to fitness costs of chromosomal replacement, and an important goal for future work will be to understand the fitness consequences of chromosomal replacement in other dominant hybrid pathogens.

## Acknowledgements

This project was funded by a Wellcome Trust Grant (106918/Z/15/Z), held by RCM. JLG was supported by funding from the Biotechnology and Biological Sciences Research Council (BBSRC) (BB/M011224/1). DJW is a Sir Henry Dale Fellow, jointly funded by the Wellcome Trust and the Royal Society (101237/Z/13/B). This work was funded by a Big Data Institute Robertson Fellowship (DJW). We thank the Oxford Genomics Center (funded by Wellcome Trust Grant 203141/Z/16/Z) for the generation and initial processing of Illumina sequence data.

## Methods

### Genome data retrieval and processing

Sequences with an MLST profile corresponding to ST239 were identified within the Staphopia database^43^. The isolation date and geographic location were extracted from the metadata files. Additional ST239 sequences with corresponding isolation date and location metadata were identified from the NCBI database using MegaBLAST^71^ and through a literature search. A total of 96 ST239 sequences were selected and downloaded from EMBL-EBI (Supplementary Table 1A). Similarly, a total of 57 ST30 and 111 ST8 genomes were identified and downloaded from EMBL-EBI (Supplementary Table 1B and 1C).

Where the sequences were downloaded as assemblies or complete genomes, raw sequence reads were simulated with dwgsim v0.1.11^72^. Illumina reads were trimmed using Trimmomatic^73^, and bwa v0.7.15 and SAMtools v1.3.1^74^ were used to map the ST239 sequences to the ST239 reference sequence (accession number FN433596). ST30 sequences were mapped to both the ST239 and ST30 reference sequences (accession number LN626917); similarly, the ST8 sequences were mapped to the ST239 and ST8 reference sequences (accession number CP007690).

The boundaries of the SCC*mec*-III element were identified in the ST239 reference genome, using BLAST against the reference SCC*mec*-III element with the accession number AB037671^52^. The boundaries were confirmed using SCC*mec*Finder^75^.

The SCC*mec* type of all 2,979 ST239 sequences in the Staphopia database was predicted using the Staphopia API. The Staphopia database was also mined for all sequences predicted to contain SCC*mec*-III. The MLST of these sequences was identified using PubMLST^76^. The MLST was confirmed using ARIBA v2.13.3^77^ and the SCC*mec* type was confirmed using SCC*mec*Finder.

### Construction of ST239 phylogeny

RaxML v8.2.9^78^ was used to construct a maximum likelihood phylogeny from the collection of 96 ST239 genomes that had been mapped to the ST239 reference sequence, using a GTR model with gamma correction for among site rate variation, which was replicated for 100 bootstraps, with recombination masked using Gubbins^39^. The ST8 reference sequence, mapped to the ST239 reference sequence, was used as an out-group. Genes were annotated with Prokka v1.13^79^. Gene function for genes that could be annotated by Prokka was identified in UniProt, and MegaBLAST^71^ was used to identify genes where no annotation was found with Prokka. The pangenome and core genome (genes shared by >99% of the ST239 isolates) was extracted using Roary v3.12.0^80^.

### Estimation of time to the MRCA of the ST239 collection

The time to the MRCA for the collection of 96 ST239 genomes was initially estimated from the maximum likelihood phylogeny (with recombination masked using Gubbins^39^), using the BactDating R package linear regression function^81^. Mixed gamma, strict gamma and relaxed gamma evolutionary models were run for 10,000,000 MCMC steps to estimate a time to the MRCA of the ST239 whole-genome sequences. The Effective Sample Size (ESS) values of all parameters were greater than 100, indicating adequate sampling of the posterior distribution.

A total of 6,819 variant sites were extracted from the ST239 sequence alignments, after recombination was masked using Gubbins^39^, using snp-sites v2.3.3^82^. Bayesian phylogenetic analysis was also carried out using BEAST version 1.10.4^38^, using the GTR nucleotide substitution model with all combinations of the strict and uncorrelated relaxed molecular clock models, and constant and exponential growth models. The XML file was edited to reflect the number of unchanging sites in the original alignments. For each model, three independent MCMC chains with 300,000,000 steps were run and combined, with path sampling/stepping-stone sampling every 100 steps. In all cases, the Bayes Factor showed no significant difference in the likelihood of the different models, and therefore the estimated time to the MRCA from the simplest model (strict molecular clock, constant population size) was recorded. The burn-in was set at 10%, and runs were combined using LogCombiner, with a re-sample size of 10,000. The MRCA and evolutionary rates were estimated with 95% HPD intervals.

BEAST analysis was repeated for 95 ST239 SCC*mec*-III element sequences (*N =* 54 variant sites), using the same clock and nucleotide substitution models as previous (one ST239 sequence was removed from the analysis due to low mapping quality of the SCC*mec*-III region).

### BLAST comparison of the ST239, ST30 and ST8 core genes

A multiple sequence alignment was constructed from the 96 ST239, 57 ST30 and 111 ST8 genomes that had been mapped to the ST239 reference genome. The shared pan-genome was calculated (*N* = 2,962 genes), and the core genes that were shared between >99% of all 264 genomes were extracted using Roary v3.12.0^77^ (*N* = 1,980 core genes). EMBOSS^83^ was used to generate ST-specific consensus sequences from the core genes of each ST (ST239, ST8 and ST30). For each gene in the ST239 consensus sequence, the percentage identities compared to the homologous gene in the ST30 and ST8 consensus sequences were calculated using MegaBLAST^71^. The ST239 core genes were defined as “acquired” (i.e. from the ST30-like region) or “backbone” (i.e. from the ST8-like region) depending on their position in the genome and similarity to the ST30 and ST8 consensus sequences.

### Identifying the closest known ancestor of the acquired and backbone regions

The consensus sequences of the 316 core genes from the acquired region of the 96 ST239 sequences were queried for similar sequences in BIGSI using a k-mer threshold of 99. This allowed for an average divergence of around 10 SNPs per gene. The most closely related sequences were identified by ST using the Staphopia API. The MLST was double checked using ARIBA, and 28 sequences were removed due to uncertainty in typing, which indicated contamination.

The consensus sequences of the 1,659 core genes from the backbone region of the 96 ST239 sequences were also queried for similar sequences in BIGSI using a k-mer threshold of 99, and Staphopia, as above. This allowed for an average divergence of around 9 SNPs per gene.

Sequences that shared at least 99% k-mer identity with over 200 of the ST239 core genes from the acquired region, and shared two or fewer MLST alleles with ST239, were downloaded from the EBI database. Only a selection of the twelve most closely related ST30, ST36 and ST39 sequences were included. These 143 sequences were mapped to the ST239 TW20 reference sequence, as above. Four sequences were removed due to poor mapping quality (>25% gaps). The acquired region was extracted (minus the SCC*mec* element) and combined into a multifasta alignment with the acquired regions from the 96 ST239 sequences and 57 ST30 sequences that were described previously.

Sequences that shared at least 99% k-mer identity with over 1,400 of the ST239 core genes from the backbone region, and were not previously identified as ST239-like, were then downloaded from the EBI database. All 32 non-ST239-like sequences were identified using the Staphopia API as ST8. These 32 ST8 sequences were mapped to the ST239 TW20 reference sequence, as above. The backbone region was extracted and combined into a multifasta alignment with the backbone regions from the 96 ST239 sequences and 111 ST8 sequences that were described previously.

All variant sites were extracted using snp-sites v2.3.3, and RaxML was used to estimate a maximum-likelihood phylogeny of all 292 acquired-region sequences, using a GTR model with gamma correction for among site rate variation and replicated for 100 bootstraps, after recombination was masked using Gubbins as previous. This was outgroup-rooted to the corresponding region of the ST8 reference sequence. A maximum-likelihood phylogeny was also estimated for the backbone region sequences, which was outgroup-rooted to the corresponding region of the ST30 reference sequence.

The acquired region of the 96 ST239 sequences and the closest related non-ST239 clade, consisting of six ST30 isolates, were combined into a multiple sequence alignment. Variant sites were extracted using snp-sites v2.3.3. To calculate the time to the MRCA, Bayesian phylogenetic analysis was carried out using BEAST version 1.10.4, as previous (for the GTR nucleotide substitution model with all combinations of the strict and uncorrelated relaxed molecular clock models, and constant and exponential growth models). This was repeated for the backbone region of the 96 ST239 sequences and the closest related non-ST239 clade, consisting of 18 ST8 isolates (one ST8 sequence was removed from the analysis, as it had no associated date of isolation). In both cases, the Bayes Factor showed no significant difference in the likelihood of the different models, and therefore the estimated time to the MRCA from the simplest model (strict molecular clock, constant population size) was recorded.

### Visualisation of recombination in ST239

ClonalFrameML v1.0-20^44^ was used to estimate regions of recombination from the ST239 phylogeny, before recombination was masked. Ten isolates from the TW20-like clade (Supplementary Table 5) were excluded from this analysis due to high sequence similarity to the ST239 reference sequence.

### Estimating evolutionary rates of the acquired and backbone regions

The evolutionary rates of the acquired and backbone regions of ST239 were estimated using BEAST analysis on 5,353 variant sites from the 96 ST239 backbone region sequences, and 1,449 variant sites from the ST239 acquired region sequences. The GTR nucleotide substitution model was used with all combinations of the strict and uncorrelated relaxed molecular clock models, and constant and exponential growth models, as previous.

### McDonald-Kreitman comparison of ST239, ST30 and ST8 core genes

All 1,980 core genes from the 96 ST239 genomes, 111 ST8 genomes and 57 ST30 genomes were converted into amino acid sequence alignments. Each of the ST239, ST30 and ST8 consensus core gene sequences that were generated previously using EMBOSS^80^ were also converted into consensus amino acid sequences.

The number of fixed synonymous and fixed non-synonymous SNPs was calculated for core genes within the whole ST239 genome, core genes within the backbone region, and core genes within the acquired region using snp-sites v2.3.3. This analysis was repeated for the 111 ST8 genomes collection and the 57 ST30 genomes collection. The McDonald-Kreitman neutrality index (N) was calculated as N = (P_n_/P_s_)/(D_n_/D_s_) where N is the net neutrality index, P_n_ is the number of non-synonymous polymorphisms, P_s_ is the number of synonymous polymorphisms, D_n_ is the number of non-synonymous substitutions, and D_s_ is the number of synonymous substitutions.

### Competition experiments

Cryostocks of the fifteen isolates were streaked on TSA, and incubated at 37°C for 24 hours. Single colonies were incubated for 24 hours in 3 mL TSB at 37°C with 225 RPM shaking. One mL of each culture was combined into a single mixture and mixed thoroughly. Genomic DNA was extracted and purified from 1 mL of the mixture, and sequencing was carried out, as previous.

Six mL of the mixture was pelleted and washed three times in PBS, and separated into six 1 mL aliquots. These were diluted 50x in either TSB, BHI or PS. A total of 3 mL of each mixed culture was incubated at 37°C with 225 RPM shaking for 24 hours. After 24 hours, genomic DNA was extracted and purified, and sequencing was carried out, as previous. All competition experiments were repeated in triplicate.

The DNA sequences from before and after each competition were mapped to the consensus sequence, as previous. The number of reads supporting each unique variant site allele (for both the reference allele and the variant allele) was determined from the mapping of the raw sequence reads from each competition experiment. Any sites that were supported with a total of three reads or less, for both the variant allele and the consensus allele, were removed from the analysis, to reduce the number of incorrect alleles due to sequencing error.

For each competition experiment, the number of supporting reads for each unique variant site was recorded. From this, the average coverage of all variants that were unique to each isolate was determined, before and after being exposed to the competition conditions for 24 hours, to determine how the proportion of each isolate changed during each competition. The limit of detection for each isolate was also calculated, as *M*_Z_=4/((*N*_Z_ *+ N*_z_’)/*V*), where *M* is the minimum detection limit for isolate *Z*, *N*_Z_ is the total number of reads in support of isolate *Z* at sites unique to isolate *Z*, *N*_Z_’ is the total number of reads in support of non-*Z* isolates at sites unique to isolate *Z*, and *V* is the number of variant sites that are unique to isolate *Z*. Any isolate with an average coverage that was lower than the limit of detection was called at the limit of detection for that isolate. Raw competitive ability was calculated as the difference in log relative abundance of each isolate before and after competition in each replicate. Raw competition values were then re-scaled, such that the mean competitive ability in each culture medium was equal to zero. This small correction factor was used to account for the fact that the true final density of some isolates was below the minimal detection threshold.

### Sequencing DNA from isolates for competition experiments

Cryostocks of the fifteen isolates were streaked on TSA, and incubated at 37°C for 24 hours. Single colonies were incubated for 24 hours in 3 mL TSB at 37°C with 225 RPM shaking. Genomic DNA was extracted and purified using the Qiagen DNeasy Blood and Tissue kit, and following the protocol for purification of bacterial or yeast DNA with enzymatic lysis, using QIAcube. Sequencing was carried out using an Illumina HiSeq4000 system with 150 bp paired-end reads by the Oxford Genomic Centre, Wellcome Trust Centre for Human Genetics, University of Oxford, UK.

The sequences were *de novo* assembled into contigs using SPAdes v3.13.0^84^, and ordered against the ST239, ST30 or ST8 reference sequences using abacas v1.3.1^85^. Sequences were annotated using PROKKA or UniProt, as previous. Using Roary, core genes were defined as genes that were shared between all 15 isolates. Unique variant alleles were defined for each isolate, which were identified in only a single isolate, using snp-sites v2.3.379.

To verify the validity of each unique variant site, each DNA sequence was re-mapped to the consensus sequence using bwa and SAMtools^71^. The number of reads supporting each unique variant site was extracted, and only unique variant sites that could be used to accurately identify a single isolate and that were supported with 4 or more reads were included as true unique variant sites.

### AMR profiles

Cryostocks of the fifteen isolates were streaked on TSA, and incubated at 37°C for 24 hours. Single colonies were incubated for 24 hours in 3 mL MH2 at 37°C with 225 RPM shaking. Cultures were diluted 100x in MH2 broth (to a density of ~1 ×10^6^ CFUs/mL), according to the MicroNaut evaluation protocol for MicroNaut-S MRSA/GP. 100 VL was added to each well of a MicroNaut-S MRSA/GP plate and incubated for 18 hours at 37°C with 225 RPM shaking, after which OD_595_ was measured. AMR breakpoints were assessed according to EUCAST^86^. This was repeated in triplicate for each isolate. Tests for ampicillin and penicillin always gave the same results, hence we considered the result from these two antibiotics as a single test score.

