## Supplementary Information for "Evolutionary processes driving the rise and fall of *Staphylococcus aureus* ST239, a dominant hybrid pathogen"

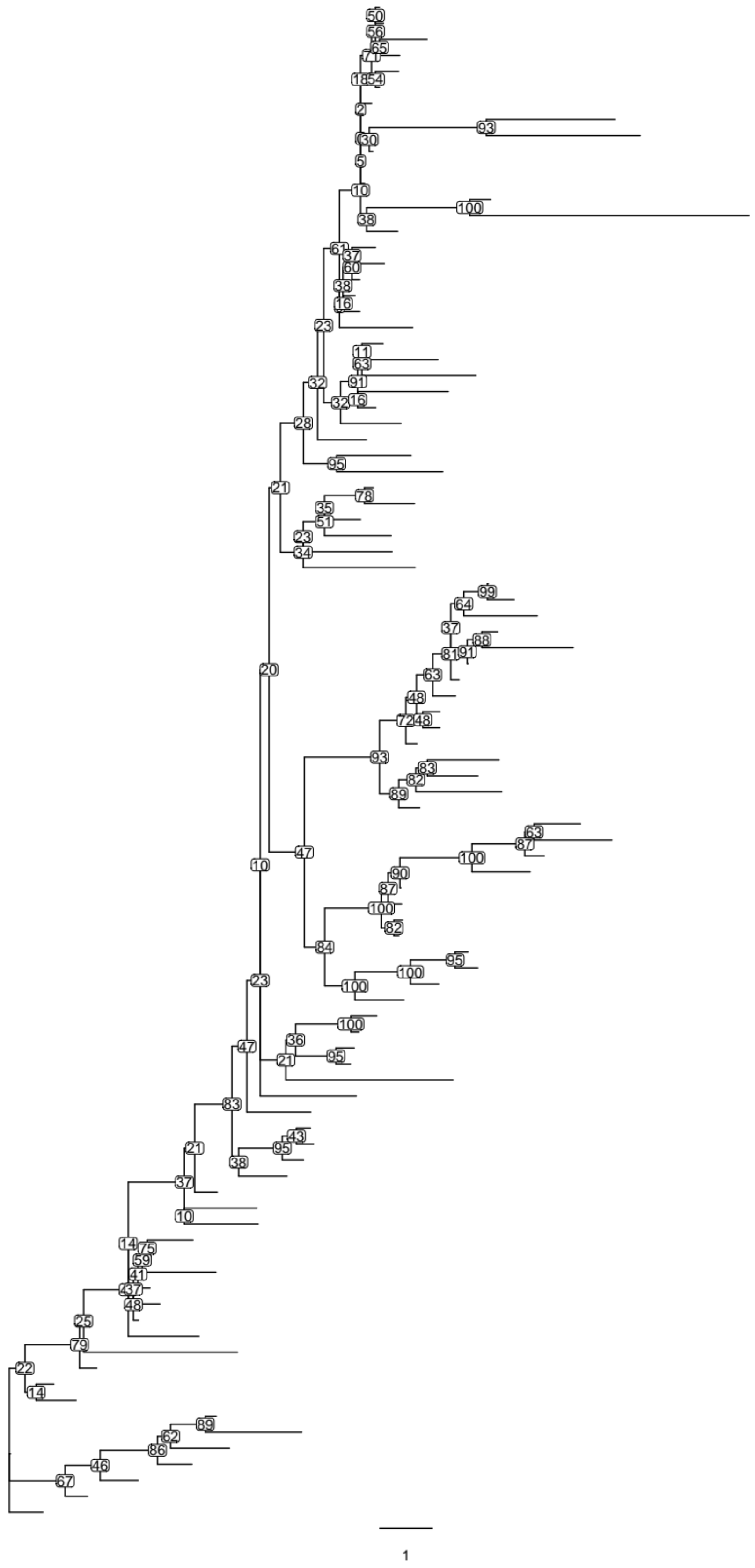


**Supplementary Figure 1.** Midpoint rooted maximum-likelihood phylogeny of the SCC*mec*-III region of ST239. Units in total number of SNPs.

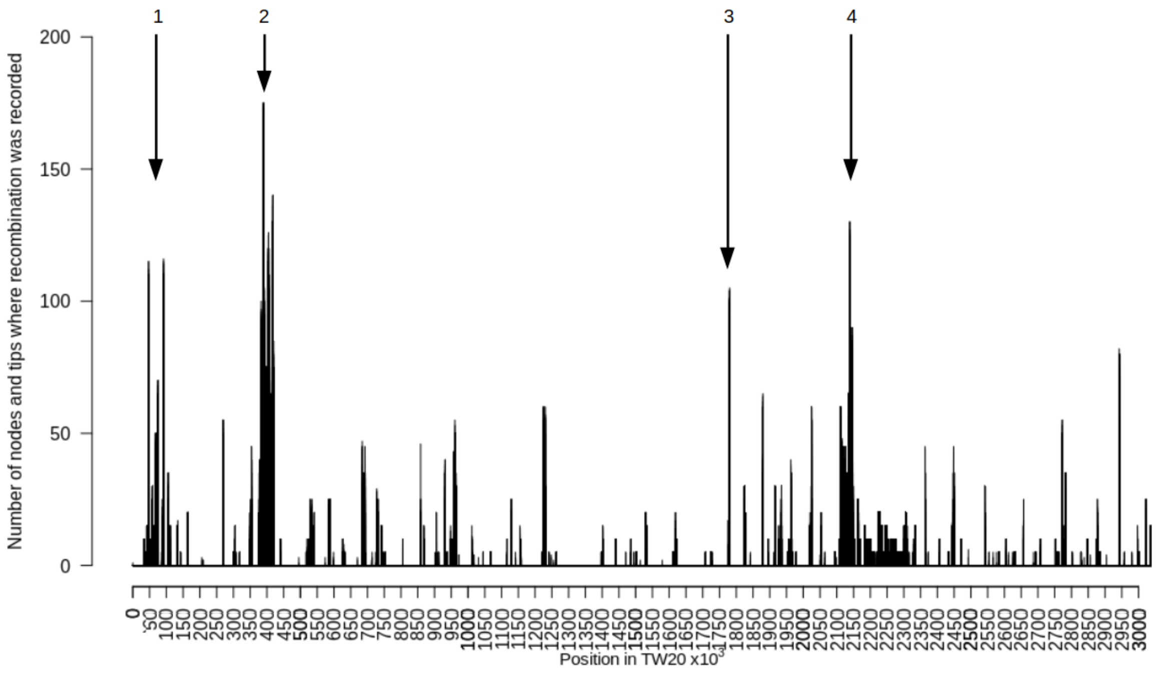


**Supplementary Figure 2**. Number of nodes and tips affected by recombination, as estimated by ClonalFrameML at each position along the TW20 genome. Regions of high recombination are labelled 1, 2, 3 and 4.

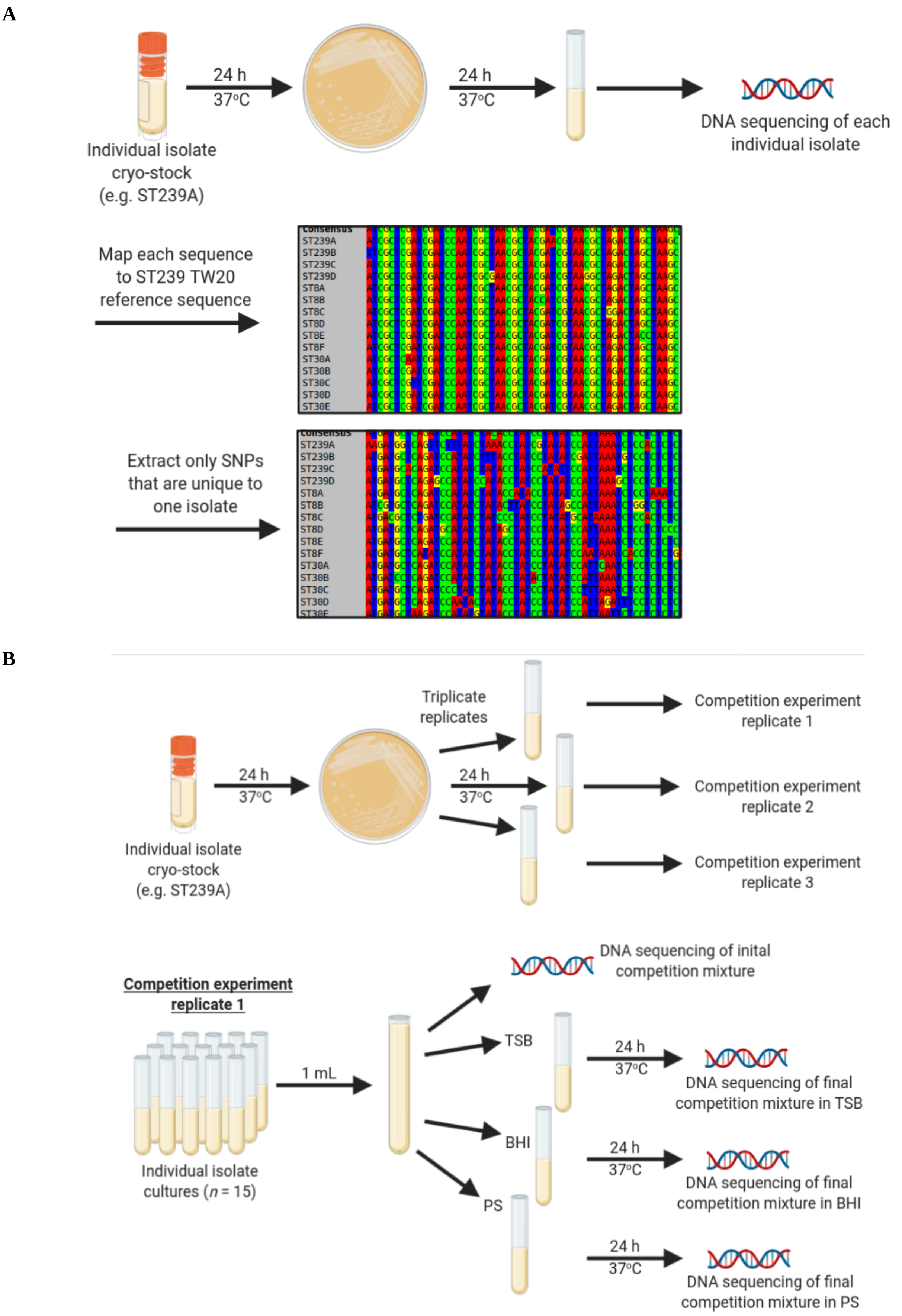
A
24 h
37°C
24 h
37°C
DNA sequencing of each individual isolate
Individual isolate cryo-stock (e.g. ST239A)
Map each sequence to ST239 TW20 reference sequence
Consensus
ST239A
ST239B
ST239C
ST239D
ST8A
ST8B
ST8C
ST8D
ST8E
ST8F
ST30A
ST30B
ST30C
ST30D
ST30E
Extract only SNPs that are unique to one isolate
B
Triplicate replicates
Competition experiment replicate 1
24 h
37°C
24 h
37°C
Competition experiment replicate 2
Competition experiment replicate 3
Individual isolate cryo-stock (e.g. ST239A)
DNA sequencing of inital competition mixture
Competition experiment replicate 1
TSB
1 mL
24 h
37°C
DNA sequencing of final competition mixture in TSB
BHI
24 h
37°C
DNA sequencing of final competition mixture in BHI
PS
24 h
37°C
DNA sequencing of final competition mixture in PS
Individual isolate cultures (n = 15)

**C**

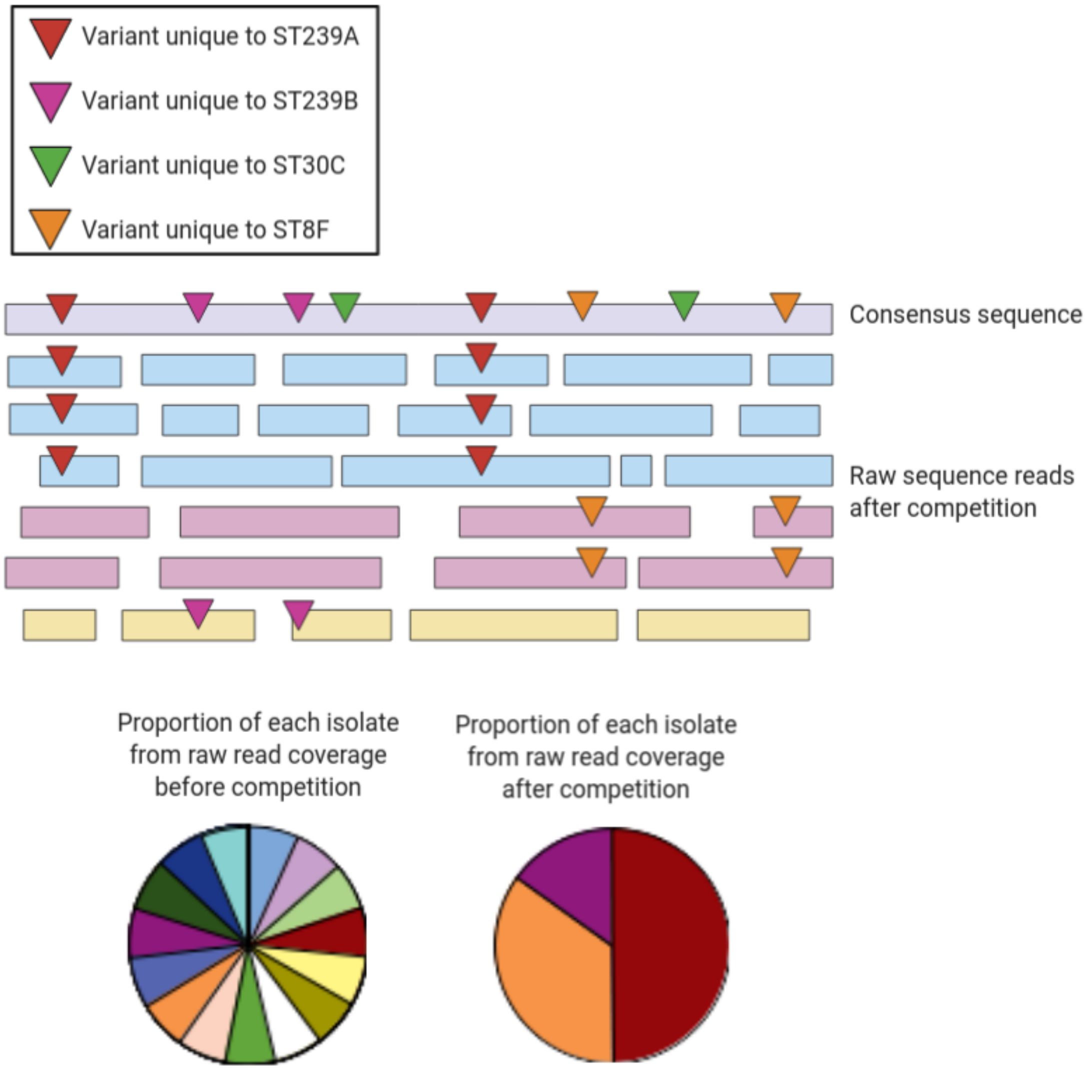


**Supplementary Figure 3A.** Schematic of unique variant site identification protocol. **3B.** Schematic of competition experimental protocol. **3C.** Schematic of protocol to estimate competitive ability.

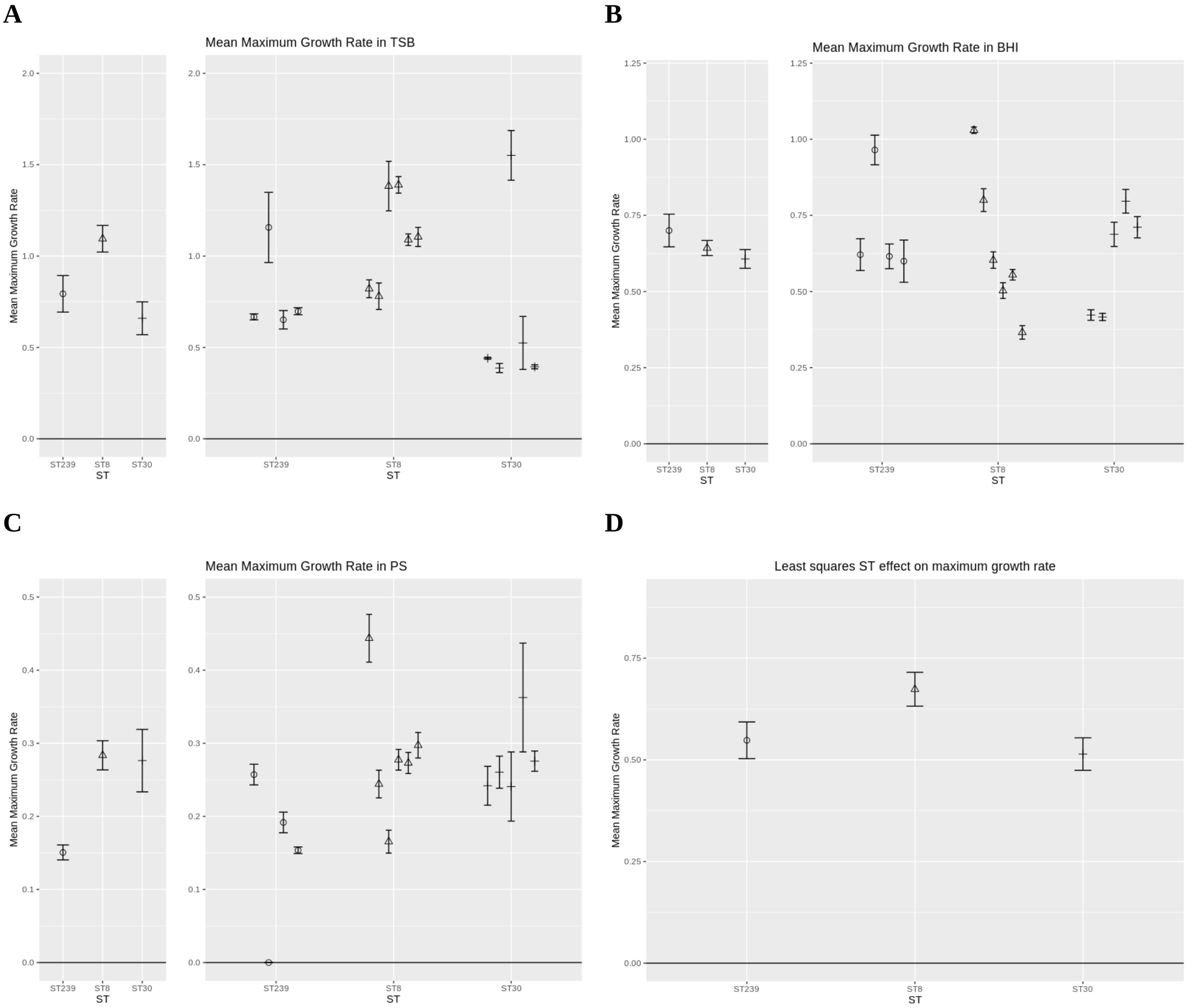


**Supplementary Figure 4.** Growth rates (+/- s.e.; $N$ = 9) of ST239, ST8 and ST30 isolates in TSB (A), BHI (B), PS (C). Panel D shows the mean (+/- s.e) growth rate of each ST across all three media. Both ST239 and ST30 have reduced growth rate relative to ST8 (ANOVA; $P$ <0.05). Cryostocks of the fifteen isolates were streaked on TSA, and incubated at 37$^{O}$C for 24 hours. Single colonies were incubated for 24 hours in 3 mL TSB at 37$^{O}$C with 225 RPM shaking. 1 mL of each culture was pelleted and washed three times in PBS, then diluted 50x in either TSB, BHI or PS. 100 μL of diluted culture was added to each well of a 96 well plate, in triplicate, and incubated at 37$^{O}$C with 225 RPM shaking. $OD_{595}$ was measured every 20 min over 24 hours, or until growth rate had reached plateau. This was repeated five times for each isolate. Growth curves were generated and analysed with Growthcurver[i] to calculate an exponential growth rate for each culture.

| Sequence ID | GenBank Accession Number (Assembly) or Run Number (SRA) | Country of isolation | Continent of isolation | Date of isolation | Sequence data format | SNP divergence from ST239 reference | PubMed ID |
|---|---|---|---|---|---|---|---|
| CTXC01 | GCA_001230325.1 | Argentina | South America | 1996 | Assembly | 5814 | 23270620 |
| CTXW01 | GCA_001229045.1 | Argentina | South America | 1996 | Assembly | 5418 | 23270620 |
| CTXX01 | GCA_001226505.1 | Argentina | South America | 1996 | Assembly | 4785 | 23270620 |
| AGT120 | ERR064919 | Argentina | South America | 1998 | Short read | 3760 | NA |
| ERS640727 | ERR732861 | Australia | Oceania | 1980 | Short read | 2710 | 25736880 |
| ERS640783 | ERR732875 | Australia | Oceania | 1982 | Short read | 2689 | 25736880 |
| ERS640788 | ERR732883 | Australia | Oceania | 1998 | Short read | 4834 | 25736880 |
| ERS640754 | ERR732893 | Australia | Oceania | 2002 | Short read | 4084 | 25736880 |
| ERS640785 | ERR732882 | Australia | Oceania | 2012 | Short read | 658 | 25736880 |
| CP005288 | GCA_000418345.1 | Brazil | South America | 1993 | Assembly | 6981 | 23873917 |
| CP009681 | GCA_000769575.1 | Brazil | South America | 1996 | Assembly | 6435 | 27152133 |
| CP012011 | GCA_001515745.1 | Brazil | South America | 2001 | Assembly | 6471 | NA |
| LDPC01 | GCA_001677355.1 | Brazil | South America | 2002 | Assembly | 7480 | NA |
| CHL1 | ERR006512 | Chile | South America | 1997 | Short read | 3766 | 20093474 |
| CHL151 | ERR006510 | Chile | South America | 1998 | Short read | 3873 | 20093474 |
| LGWM01 | GCA_002267325.1 | Chile | South America | 2012 | Assembly | 5172 | 28760895 |
| CHI59 | ERR064937 | China | Asia | 1998 | Short read | 3260 | NA |
| CP007447 | GCA_000709475.1 | China | Asia | 2004 | Assembly | 2138 | 239544 |
| CP002643 | GCA_000204665.1 | China | Asia | 2006 | Assembly | 4334 | 21551295 |
| CP006838 | GCA_000485885.1 | China | Asia | 2010 | Assembly | 4883 | 24309740 |
| 3HK | ERR006470 | Czechia | Europe | 2000 | Short read | 4097 | 20093474 |
| 2A8 | ERR006540 | Czechia | Europe | 2001 | Short read | 3479 | 20093474 |
| DEN907 | ERR006514 | Denmark | Europe | 2001 | Short read | 665 | 20093474 |
| CTWW01 | GCA_001239465.1 | Denmark | Europe | 2006 | Assembly | 3530 | 23270620 |
| CTXO01 | GCA_001227085.1 | Denmark | Europe | 2009 | Assembly | 2966 | NA |
| CTWS01 | GCA_001229625.1 | Egypt | Africa | 2005 | Assembly | 5880 | NA |
| ERS026791 | ERR033471 | France | Europe | 2006 | Short read | 4009 | NA |
| D8 | ERR1213802 | Gambia | Africa | 2007 | Short read | 1071 | 27474712 |
| GRE108 | ERR064936 | Greece | Europe | 1998 | Short read | 60322 | NA |
| GRE18 | ERR064902 | Greece | Europe | 1998 | Short read | 4830 | NA |
| GRE4 | ERR064904 | Greece | Europe | 1998 | Short read | 5042 | NA |
| GRE317 | ERR064903 | Greece | Europe | 1999 | Short read | 4993 | NA |
| HUSA304 | ERR064928 | Hungary | Europe | 1993 | Short read | 3699 | NA |
| HU106 | ERR064927 | Hungary | Europe | 1996 | Short read | 3712 | NA |
| HU109 | ERR064909 | Hungary | Europe | 1996 | Short | 5122 | NA |

| | | | | | | | |
|---|---|---|---|---|---|---|---|
| | | | | | read | | |
| HUR18 | ERR064910 | Hungary | Europe | 1997 | Short read | 4666 | NA |
| CTXS01 | GCA_001224885.1 | India | Asia | 2006 | Assembly | 6834 | NA |
| LWAK01 | GCA_001652195.1 | India | Asia | 2014 | Assembly | 6279 | NA |
| LWAM01 | GCA_001652225.1 | India | Asia | 2014 | Assembly | 6648 | NA |
| NESW01 | GCA_002738565.1 | India | Asia | 2014 | Assembly | 11676 | NA |
| MLQB01 | GCA_001855085.1 | India | Asia | 2015 | Assembly | 6528 | NA |
| CTXR01 | GCA_001232225.1 | Lithuania | Europe | 1996 | Assembly | 5180 | 23270620 |
| CTYA01 | GCA_001239185.1 | Lithuania | Europe | 1996 | Assembly | 5444 | 23270620 |
| CTYC01 | GCA_001226905.1 | Lithuania | Europe | 1996 | Assembly | 5438 | 23270620 |
| CTWZ01 | GCA_001237525.1 | Malaysia | Asia | 1996 | Assembly | 1629 | 23270620 |
| JQNU01 | GCA_000787025.1 | Malaysia | Asia | 2003 | Assembly | 2310 | NA |
| AMRE01 | GCA_000313005.1 | Malaysia | Asia | 2009 | Assembly | 1899 | 23405328 |
| ANPO01 | GCA_000784385.1 | Malaysia | Asia | 2009 | Assembly | 2127 | 25197474 |
| JKD6009 | ERR732894 | New Zealand | Oceania | 2003 | Short read | 4714 | 25736880 |
| CP002120 | GCA_000145595.1 | New Zealand | Oceania | 2013 | Assembly | 5811 | 20802046 |
| LGWY01 | GCA_002267125.1 | Peru | South America | 2011 | Assembly | 6284 | 28760895 |
| LGWZ01 | GCA_002267145.1 | Peru | South America | 2011 | Assembly | 5747 | 28760895 |
| MIHO01 | GCA_002260265.1 | Peru | South America | 2013 | Assembly | 6203 | 28760895 |
| CTXV01 | GCA_001226265.1 | Poland | Europe | 1996 | Assembly | 7264 | 23270620 |
| ERS026930 | ERR038683 | Poland | Europe | 2006 | Short read | 4873 | NA |
| ERS049893 | ERR064932 | Portugal | Europe | 1990 | Short read | 3125 | NA |
| ICP5014 | ERR064934 | Portugal | Europe | 1993 | Short read | 3433 | NA |
| HSJ216 | ERR064920 | Portugal | Europe | 1998 | Short read | 3798 | NA |
| HGSA142 | ERR064922 | Portugal | Europe | 2003 | Short read | 4168 | NA |
| CTXP01 | GCA_001227985.1 | Portugal | Europe | 2005 | Assembly | 4737 | 23270620 |
| CTXK01 | GCA_001232765.1 | Romania | Europe | 2007 | Assembly | 6710 | 23270620 |
| SRS1497005 | SRR3658375 | Russia | Europe | 2011 | Short read | 4700 | 31329022 |
| SRS1496939 | SRR3658370 | Russia | Europe | 2013 | Short read | 4154 | 31329022 |
| SRS2490451 | SRR6003859 | Russia | Europe | 2016 | Short read | 4280 | 31329022 |
| ERR053014 | ERR053014 | Singapore | Asia | 1982 | Short read | 1252 | 25903077 |
| ERR053016 | ERR053016 | Singapore | Asia | 1985 | Short read | 310 | 25903077 |
| ERR053031 | ERR053031 | Singapore | Asia | 1997 | Short read | 370 | 25903077 |
| ERR030243 | ERR030243 | Singapore | Asia | 2002 | Short read | 499 | 25903077 |
| ERR029473 | ERR029473 | Singapore | Asia | 2009 | Short read | 334 | 25903077 |
| LFED02 | GCA_001184285.2 | South Korea | Asia | 2014 | Assembly | 994 | 27313284 |
| CP013957 | GCA_001641025.1 | South Korea | Asia | 2011 | Assembly | 1092 | NA |
| CTXI01 | GCA_001233925.1 | Spain | Europe | 1996 | Assembly | 6603 | 23270620 |
| AHZL01 | GCA_000401475.1 | Switzerland | Europe | 1990 | Assembly | 11802 | NA |
| ERR064946 | ERR064946 | Thailand | Asia | 2006 | Short read | 948 | NA |
| CTWU01 | GCA_001239585.1 | Thailand | Asia | 2007 | Assembly | 4380 | 23270620 |
| ERR023864 | ERR023864 | Thailand | Asia | 2008 | Short read | 1143 | 25491771 |
| BPH0365 | ERR732813 | Turkey | Europe | 1996 | Short read | 3329 | 25736880 |
| CTWE01 | GCA_001225725.1 | Turkey | Europe | 2006 | Assembly | 6223 | 23270620 |

| | | | | | | | |
|---|---|---|---|---|---|---|---|
| CP012015 | GCA_001515665.1 | Turkey | Europe | 2007 | Assembly | 6360 | 23270620 |
| CTWD01 | GCA_001226925.1 | Turkey | Europe | 2007 | Assembly | 6328 | 23270620 |
| CTWI01 | GCA_001236045.1 | Turkey | Europe | 2007 | Assembly | 6148 | 23270620 |
| CTYZ01 | GCA_001234465.1 | Turkey | Europe | 2008 | Assembly | 6286 | 23270620 |
| CTWH01 | GCA_001226645.1 | Turkey | Europe | 2009 | Assembly | 6457 | 23270620 |
| FN433596 | GCA_000027045.1 | UK | Europe | 2003 | Assembly | 0 | 19948800 |
| ERR1212592 | ERR1212592 | UK | Europe | 2009 | Short read | 2029 | 28348843 |
| FFPZ01 | GCA_900043625.1 | UK | Europe | 2010 | Assembly | 648 | 26672018 |
| FGZX01 | GCA_900058615.1 | UK | Europe | 2010 | Assembly | 1194 | 26672018 |
| SRS1974492 | SRR5251374 | UK | Europe | 2011 | Short read | 4477 | 29256859 |
| URU34 | ERR064925 | Uruguay | South America | 1997 | Short read | 3529 | NA |
| URU110 | ERR006556 | Uruguay | South America | 1998 | Short read | 3047 | NA |
| JXZH02 | GCA_001018635.2 | USA | North America | 1981 | Assembly | 4795 | 31346170 |
| R35 | ERR006550 | USA | North America | 1987 | Short read | 4102 | NA |
| LHH1 | ERR006548 | USA | North America | 1994 | Short read | 4436 | NA |
| BK2421 | ERR064899 | USA | North America | 1996 | Short read | 4027 | NA |
| SR2852 | SRR1177896 | USA | North America | 2005 | Short read | 754 | 24787619 |
| CTXQ01 | GCA_001234765.1 | Vietnam | Asia | 2004 | Assembly | 1949 | 23270620 |

**Supplementary Table 1A**. Metadata of the ST239 sequences used in this study.

| **Sequence ID** | **GenBank Accession Number (Assembly) or Run Number (SRA)** | **Country of isolation** | **Continent of isolation** | **Date of isolation** | **Sequence data format** |
|---|---|---|---|---|---|
| ERR1535436 | ERR1535436 | Australia | Oceania | 1989 | Short read |
| ERR1535439 | ERR1535439 | Australia | Oceania | 2003 | Short read |
| ERR1535431 | ERR1535431 | Australia | Oceania | 2008 | Short read |
| ERR1535432 | ERR1535432 | Australia | Oceania | 2010 | Short read |
| ERR1535433 | ERR1535433 | Australia | Oceania | 2011 | Short read |
| CP007690 | GCA_000695875.1 | Belgium | Europe | 1905 | Assembly |
| ERR033306 | ERR033306 | Belgium | Europe | 2006 | Short read |
| ERR033347 | ERR033347 | Belgium | Europe | 2006 | Short read |
| CP019590 | GCA_002088995.1 | Canada | North America | 2004 | Assembly |
| ERR381019 | ERR381019 | Canada | North America | 2008 | Short read |
| ERR380998 | ERR380998 | Canada | North America | 2009 | Short read |
| ERR380994 | ERR380994 | Canada | North America | 2010 | Short read |
| ERR381026 | ERR381026 | Canada | North America | 2011 | Short read |
| CP007676 | GCA_001045995.2 | Colombia | South America | 2006 | Assembly |
| CP007672 | GCA_001045795.2 | Colombia | South America | 2007 | Assembly |
| CP007674 | GCA_001021895.1 | Colombia | South America | 2007 | Assembly |

| | | | | | |
|---|---|---|---|---|---|
| CP027101 | GCA_001471715.2 | Colombia | South America | 2014 | Assembly |
| ERR1588671 | ERR1588671 | Denmark | Europe | 1957 | Short read |
| ERR1588666 | ERR1588666 | Denmark | Europe | 1968 | Short read |
| ERR1712346 | ERR1712346 | Denmark | Europe | 2004 | Short read |
| ERR1712341 | ERR1712341 | Denmark | Europe | 2007 | Short read |
| SRR6169019 | SRR6169019 | Denmark | Europe | 2016 | Short read |
| SRR1955591 | SRR1955591 | France | Europe | 1999 | Short read |
| ERR033450 | ERR033450 | France | Europe | 2006 | Short read |
| ERR033462 | ERR033462 | France | Europe | 2006 | Short read |
| CP026961 | GCA_001018725.2 | French Guyana | South America | 1996 | Assembly |
| ERR1535412 | ERR1535412 | Gabon | Africa | 2005 | Short read |
| ERR1535387 | ERR1535387 | Gabon | Africa | 2009 | Short read |
| ERR1535352 | ERR1535352 | Gabon | Africa | 2011 | Short read |
| ERR1143478 | ERR1143478 | Gabon | Africa | 2012 | Short read |
| ERR1535423 | ERR1535423 | Gabon | Africa | 2013 | Short read |
| SRR4359408 | SRR4359408 | Germany | Europe | 2001 | Short read |
| CP003033 | GCA_000245495.1 | Germany | Europe | 2002 | Assembly |
| ERR1535365 | ERR1535365 | Germany | Europe | 2006 | Short read |
| ERR593213 | ERR593213 | Germany | Europe | 2013 | Short read |
| CP022290 | GCA_002786465.1 | Germany | Europe | 2014 | Assembly |
| ERR1535441 | ERR1535441 | Ghana | Africa | 2005 | Short read |
| ERR1535358 | ERR1535358 | Ghana | Africa | 2011 | Short read |
| ERR107788 | ERR107788 | Ireland | Europe | 2001 | Short read |
| ERR118522 | ERR118522 | Ireland | Europe | 2001 | Short read |
| ERR124443 | ERR124443 | Ireland | Europe | 2004 | Short read |
| ERR211899 | ERR211899 | Ireland | Europe | 2004 | Short read |
| ERR124475 | ERR124475 | Ireland | Europe | 2010 | Short read |
| ERR033701 | ERR033701 | Italy | Europe | 2006 | Short read |
| AP014921 | GCA_002356675.1 | Japan | Asia | 2013 | Assembly |
| SRR917579 | SRR917579 | Luxembourg | Europe | 1997 | Short read |
| SRR917586 | SRR917586 | Luxembourg | Europe | 2005 | Short read |
| ERR1535363 | ERR1535363 | Luxembourg | Europe | 2006 | Short read |
| SRR917585 | SRR917585 | Luxembourg | Europe | 2007 | Short read |
| SRR917652 | SRR917652 | Luxembourg | Europe | 2010 | Short read |
| ERR1143491 | ERR1143491 | Mozambique | Africa | 2012 | Short read |
| SRR3658373 | SRR3658373 | Russia | Europe | 2013 | Short read |
| SRR5100325 | SRR5100325 | Russia | Europe | 2016 | Short read |
| SRR5100330 | SRR5100330 | Russia | Europe | 2016 | Short read |
| ERR033296 | ERR033296 | Spain | Europe | 2006 | Short read |
| CP014362 | GCA_003595365.1 | Suriname | South America | 2013 | Assembly |
| CP014384 | GCA_003595485.1 | Suriname | South America | 2013 | Assembly |
| CP014444 | GCA_002000865.1 | Suriname | South America | 2013 | Assembly |

| | | | | | |
|---|---|---|---|---|---|
| ERR033573 | ERR033573 | Sweden | Europe | 2006 | Short read |
| ERR033574 | ERR033574 | Sweden | Europe | 2006 | Short read |
| SRR1786287 | SRR1786287 | Switzerland | Europe | 2007 | Short read |
| SRR1790318 | SRR1790318 | Switzerland | Europe | 2007 | Short read |
| ERR505339 | ERR505339 | Tanzania | Africa | 2008 | Short read |
| ERR505360 | ERR505360 | Tanzania | Africa | 2008 | Short read |
| ERR1143410 | ERR1143410 | Tanzania | Africa | 2011 | Short read |
| ERR505358 | ERR505358 | Tanzania | Africa | 2013 | Short read |
| ERR1213800 | ERR1213800 | The Gambia | Africa | 2007 | Short read |
| ERR1213806 | ERR1213806 | The Gambia | Africa | 2008 | Short read |
| ERR1712350 | ERR1712350 | The Netherlands | Europe | 2005 | Short read |
| ERR1712351 | ERR1712351 | The Netherlands | Europe | 2006 | Short read |
| ERR1712352 | ERR1712352 | The Netherlands | Europe | 2007 | Short read |
| SRR5029779 | SRR5029779 | UK | Europe | 1935 | Short read |
| ERR172082 | ERR172082 | UK | Europe | 1998 | Short read |
| ERR129281 | ERR129281 | UK | Europe | 2003 | Short read |
| ERR2072416 | ERR2072416 | UK | Europe | 2012 | Short read |
| SRR5651527 | SRR5651527 | UK | Europe | 2015 | Short read |
| SRR1103475 | SRR1103475 | USA, CA | North America | 2011 | Short read |
| KK095312 | KK095312.1 | USA, CA | North America | 2012 | Assembly |
| SRR1165956 | SRR1165956 | USA, CA | North America | 2013 | Short read |
| SRR3194930 | SRR3194930 | USA, CO | North America | 2010 | Short read |
| SRR3194933 | SRR3194933 | USA, CO | North America | 2011 | Short read |
| SRR4140011 | SRR4140011 | USA, FL | North America | 2003 | Short read |
| SRR4140097 | SRR4140097 | USA, FL | North America | 2006 | Short read |
| SRR4140090 | SRR4140090 | USA, FL | North America | 2008 | Short read |
| SRR4140057 | SRR4140057 | USA, FL | North America | 2010 | Short read |
| SRR4140079 | SRR4140079 | USA, FL | North America | 2011 | Short read |
| CP016855 | GCA_001717645.3 | USA, GA | North America | 2010 | Assembly |
| CP017094 | CP017094.2 | USA, GA | North America | 2011 | Assembly |
| CP025495 | GCA_001735655.2 | USA, GA | North America | 2011 | Assembly |
| SRR1786280 | SRR1786280 | USA, IL | North America | 1965 | Short read |
| SRR497495 | SRR497495 | USA, MA | North America | 2003 | Short read |
| KB820894 | KB820894.1 | USA, MA | North America | 2007 | Assembly |
| SRR1539657 | SRR1539657 | USA, MA | North America | 2012 | Short read |
| SRR5226531 | SRR5226531 | USA, MA | North America | 2013 | Short read |
| SRR3168638 | SRR3168638 | USA, MA | North America | 2015 | Short read |
| SRR3194883 | SRR3194883 | USA, MD | North America | 2010 | Short read |
| SRR3194942 | SRR3194942 | USA, MD | North America | 2011 | Short read |
| CP020619 | GCA_002085525.1 | USA, NE | North America | 2013 | Assembly |
| ERR134641 | ERR134641 | USA, NY | North America | 2006 | Short read |
| ERR302763 | ERR302763 | USA, NY | North America | 2007 | Short read |

| | | | | | |
|---|---|---|---|---|---|
| ERR338634 | ERR338634 | USA, NY | North America | 2008 | Short read |
| ERR134699 | ERR134699 | USA, NY | North America | 2010 | Short read |
| ERR134723 | ERR134723 | USA, NY | North America | 2011 | Short read |
| SRR3194889 | SRR3194889 | USA, OR | North America | 2010 | Short read |
| SRR3194948 | SRR3194948 | USA, OR | North America | 2011 | Short read |
| SRR3194899 | SRR3194899 | USA, PA | North America | 2010 | Short read |
| SRR3194910 | SRR3194910 | USA, TN | North America | 2010 | Short read |
| SRR3194949 | SRR3194949 | USA, TN | North America | 2011 | Short read |
| CP026076 | GCA_001018685.2 | USA, TX | North America | 2002 | Assembly |
| CP013231 | GCA_001580515.1 | USA, TX | North America | 2013 | Assembly |
| SRR4019282 | SRR4019282 | USA, TX | North America | 2016 | Short read |

**Supplementary Table 1B**. Metadata of the ST8 sequences used in this study.

| **Sequence ID** | **GenBank Accession Number (Assembly) or Run Number (SRA)** | **Country of isolation** | **Continent of isolation** | **Date of isolation** | **Sequence data format** |
|---|---|---|---|---|---|
| SRR1172654 | SRR1172654 | Argentina | South America | 2013 | Short read |
| LDIT01 | GCA_001026765.1 | Brazil | South America | 2014 | Assembly |
| ERR033374 | ERR033374 | Denmark | Europe | 2006 | Short read |
| ERR1753509 | ERR1753509 | Denmark | Europe | 2013 | Short read |
| ERR1753386 | ERR1753386 | Denmark | Europe | 2015 | Short read |
| ERR1592225 | ERR1592225 | DRC | Africa | 2013 | Short read |
| ERR1592254 | ERR1592254 | DRC | Africa | 2013 | Short read |
| ERR1592169 | ERR1592169 | DRC | Africa | 2014 | Short read |
| ERR1592321 | ERR1592321 | DRC | Africa | 2015 | Short read |
| ERR033635 | ERR033635 | Finland | Europe | 2006 | Short read |
| ERR033464 | ERR033464 | France | Europe | 2006 | Short read |
| ERR038658 | ERR038658 | France | Europe | 2006 | Short read |
| SRR4359400 | SRR4359400 | Germany | Europe | 2002 | Short read |
| ERR1143353 | ERR1143353 | Germany | Europe | 2010 | Short read |
| ERR1675778 | ERR1675778 | Germany | Europe | 2014 | Short read |
| ERR1195906 | ERR1195906 | Germany | Europe | 2015 | Short read |
| ERR033537 | ERR033537 | Italy | Europe | 2006 | Short read |
| ERR033542 | ERR033542 | Italy | Europe | 2006 | Short read |
| JHEC01 | GCA_000708425.1 | Jordan | Asia | 2009 | Assembly |
| LN626917 | GCA_000953255.1 | Kenya | Africa | 2004 | Assembly |
| JHEB01 | GCA_000708405.1 | Lebanon | Asia | 2011 | Assembly |
| ERR033698 | ERR033698 | Norway | Europe | 2006 | Short read |
| ERR038674 | ERR038674 | Norway | Europe | 2006 | Short read |

| | | | | | |
|---|---|---|---|---|---|
| ERR033550 | ERR033550 | Poland | Europe | 2006 | Short read |
| ERR038685 | ERR038685 | Poland | Europe | 2006 | Short read |
| ERR038688 | ERR038688 | Poland | Europe | 2006 | Short read |
| ERR033556 | ERR033556 | Portugal | Europe | 2006 | Short read |
| LWRA01 | GCA_001639185.1 | Russia | Europe | 2013 | Assembly |
| LWRC01 | GCA_001639225.1 | Russia | Europe | 2014 | Assembly |
| LWRD01 | GCA_001639195.1 | Russia | Europe | 2014 | Assembly |
| CP009554 | GCA_000772025.1 | South Korea | Asia | 2014 | Assembly |
| ERR033291 | ERR033291 | Spain | Europe | 2006 | Short read |
| ERR033578 | ERR033578 | Sweden | Europe | 2006 | Short read |
| ERR033589 | ERR033589 | Sweden | Europe | 2006 | Short read |
| ERR033590 | ERR033590 | Sweden | Europe | 2006 | Short read |
| ERR033593 | ERR033593 | Sweden | Europe | 2006 | Short read |
| AHZJ01 | GCA_000401455.1 | Switzerland | Europe | 2005 | Assembly |
| JXHZ01 | GCA_000878025.1 | Switzerland | Europe | 2007 | Assembly |
| SRR1786285 | SRR1786285 | Switzerland | Europe | 2007 | Short read |
| ERR1143398 | ERR1143398 | Tanzania | Africa | 2011 | Short read |
| ERR1143429 | ERR1143429 | Tanzania | Africa | 2012 | Short read |
| ERR505346 | ERR505346 | Tanzania | Africa | 2012 | Short read |
| ERR505389 | ERR505389 | Tanzania | Africa | 2013 | Short read |
| ERR829972 | ERR829972 | Tanzania | Africa | 2014 | Short read |
| ERR1213763 | ERR1213763 | The Gambia | Africa | 2002 | Short read |
| ERR1213777 | ERR1213777 | The Gambia | Africa | 2004 | Short read |
| ERR1213799 | ERR1213799 | The Gambia | Africa | 2007 | Short read |
| JYAJ02 | GCA_001019455.2 | UK | Europe | 1935 | Assembly |
| ERR109495 | ERR109495 | UK | Europe | 1998 | Short read |
| ERR129280 | ERR129280 | UK | Europe | 2003 | Short read |
| SRR5251295 | SRR5251295 | UK | Europe | 2008 | Short read |
| JPWO01 | GCA_000739205.1 | UK | Europe | 2013 | Assembly |
| ERR554777 | ERR554777 | USA | North America | 2004 | Short read |
| ERR554339 | ERR554339 | USA | North America | 2009 | Short read |
| CP017684 | GCA_002633785.1 | USA | North America | 2011 | Assembly |
| SRR1159198 | SRR1159198 | USA | North America | 2012 | Short read |
| SRR5822163 | SRR5822163 | USA | North America | 2015 | Short read |

**Supplementary Table 1C**. Metadata of the ST30 sequences used in this study.

| **Substitution rate (95% HPD) ($x10^{-6}$ SNPs/site/ year)** | **Region** | **BEAST evolutionary model (95% HPD intervals)** | | | |
|---|---|---|---|---|---|
| | | **Strict clock Constant population size** | **Strict clock Exponential population size** | **Relaxed clock Constant population size** | **Relaxed clock Exponential population size** |
| | ST239 whole genome | **1.20** (1.13 - 1.28) | **1.20** (1.13 - 1.28) | **1.25** (0.99 - 1.53) | **1.29** (1.05 - 1.55) |
| | ST8-like region | **1.21** (1.13 - 1.29) | **1.21** (1.12 - 1.29) | **0.749** (0.494 - 1.08) | **1.25** (1.02 - 1.50) |
| | ST30-like region | **1.52** (1.32 - 1.71) | **1.48** (1.30 - 1.68) | **1.7** (1.31 - 2.19) | **1.63** (1.27 - 2.00) |
| **MRCA** | ST239 whole genome | **1940.1** (1934.7 - 1945.4) | **1940.1** (1934.6 - 1945.3) | **1929.1** (1899.3 - 1953.0) | **1946.5** (1931.5 - 1959.9) |
| | SCC*mec*-III | **1935.0** (1892.7 – 1967.6) | **1959.7** (1939.1 – 1976.4) | **1901.5** (1791.9 – 1968.3) | **1960.2** (1938.1 – 1977.4) |

**Supplementary Table 2A.** BEAST model estimates of time to the MRCA of ST239.

| | **BactDating evolutionary model (95% HPD)** | | |
|---|---|---|---|
| **Estimated time to MRCA** | **Mixed gamma** | **Relaxed gamma** | **Strict gamma** |
| | 1906.0 – 1949.8 | 1903.1 – 1947.3 | 1936.2 – 1943.0 |

**Supplementary Table 2B.** BactDating model estimates of time to the MRCA of ST239.

| **ST** | **Accession Number** | **Source** | **Reference** |
|---|---|---|---|
| ST22 | ERR171907 | human blood sample in England in 2011 | [ii] |
| ST5 | SRR1162455 | 1995 from a human source in Belgium | [iii] |
| ST101 | ERR086199 | Unknown | [iv] |

**Supplementary Table 3.** Non-ST239-like isolates in the Staphopia database that contain SCC*mec*-III.

| Accession Number | Date | Country | Source |
|---|---|---|---|
| SRR016399 | 1955 | UK | Unknown |
| SRR016398 | 1955 | UK | Clinical |
| SRR016780 | 1962 | Australia | Unknown |
| SRR016388 | 1965 | USA | Unknown |
| SRR016400 | 1968 | USA | Blood |
| SRR2124638 | Unknown | Unknown | Unknown |

**Supplementary Table 4A.** Accession numbers of the ST30 genomes that share the closest common ancestor with the acquired region of the ST239 genome.

| Accession Number | Date | Country | Source |
|---|---|---|---|
| ERR1588671 | 1957 | Denmark | Bacteraemia |
| ERR1588666 | 1968 | Denmark | Bacteraemia |
| ERR1712346 | 2004 | Denmark | Human, clinical |
| ERR107788 | 2001 | Ireland | Blood |
| ERR211899 | 2004 | Ireland | Blood |
| ERR124443 | 2004 | Ireland | Blood |
| ERR124475 | 2010 | Ireland | Blood |
| ERR033574 | 2007 | Sweden | Invasive human disease |
| SRR1786287 | 2007 | Switzerland (Bern) | Food |
| ERR1535423 | 2017 | Gabon | Unknown |
| ERR1213800 | 2007 | Gambia | Skin and soft tissue disease |
| ERR1143491 | 2012 | Mozambique | Human |
| ERR1535431 | 2008 | Tanzania | Nasal carriage |
| ERR505339 | 2008 | Tanzania | Nasal carriage |
| ERR505358 | 2013 | Tanzania | Nasal carriage |
| SRR1786280 | 1965 | USA (Chicago) | Human sample |
| KB820894 | 2007 | USA (Boston) | Human sample |
| ERR338634 | 2008 | USA (New York) | Soft tissue disease |
| ERR1007706 | Unknown | Unknown | Unknown |

**Supplementary Table 4B.** Accession numbers of the ST8 genomes that share the closest common ancestor with the backbone region of the ST239 genome.

| Sequence name | SNP divergence from TW20 reference |
|---|---|
| LFED02_SKorea_2014 | 994 |
| ERS640785_Australia_2012 | 658 |
| ERR030243_Singapore_2002 | 499 |
| SR2852_USA_2005 | 754 |
| ERR029473_Singapore_2009 | 334 |
| FFPZ01_UK_2010 | 648 |
| FN433596_UK_2003 | 0 |
| FGZX01_UK_2010 | 1194 |
| ERR053031_Singapore_1997 | 370 |
| ERR053016_Singapore_1985 | 310 |

**Supplementary Table 5.** ST239-like isolates that were removed from the recombination analysis using ClonalFrameML.

| Effect | df | Sum of squares | Mean squares | F-value | *p*-value |
|---|---|---|---|---|---|
| ST | 2 | 0.2605196 | 0.1302598 | 7.692 | 0.00073 |
| Media x ST | 4 | 0.5271552 | 0.1317888 | 7.7823 | 0.00001 |
| Isolate[ST] | 12 | 4.2435429 | 0.353628575 | 20.88 | $<1 \times 10^{-5}$ |
| Error | 134 | 6.9956092 | 0.0522060388 | - | - |

**Supplementary Table 6A.** Reduced ANOVA table of significant competitive ability effects, with media effect removed.

| Effect | df | Sum of squares | Mean squares | F-value | *p*-value |
|---|---|---|---|---|---|
| ST | 2 | 1.1635513 | 0.58177565 | 4.9487 | 0.0080 |
| Media | 2 | 14.238509 | 7.119255 | 145.5380 | $<1 \times 10^{-5}$ |
| Media x ST | 4 | 2.6565443 | 0.664136075 | 5.6493 | 0.0002 |
| Isolate[ST] | 12 | 3.1333904 | 0.261115867 | 2.2211 | 0.0120 |
| Error | 206(204) | 24.217540 | 0.117560874 | - | - |

**Supplementary Table 6B.** Reduced ANOVA table of significant growth rate effects, with media effect removed.

| Gene name | Genetic Region | No. SNPs | Function | General Function |
|---|---|---|---|---|
| *walK* | Acquired | 13 | Cell wall metabolism sensor histidine kinase WalK. Regulates genes involved in autolysis, biofilm formation and cell wall metabolism, and linked to vancomycin resistance. | AMR |
| *spa* | Acquired | 62 | Peptidoglycan-binding protein LysM. Virulence factor that enables host immune response evasion. Highly variable . | Virulence |
| *gsiA* | Acquired | 13 | Peptide ABC transporter ATP-binding protein. Down-regulation linked to antimicrobial peptide resistance. | AMR |
| *ausA* | Acquired | 26 | Hypothetical aureusimine synthesis protein – potential virulence factor. | Virulence |
| *manP* | Acquired | 15 | Phospho-transferase system mannose transporter subunit IIABC. | Metabolism |
| *tarL* | Acquired | 25 | Teichoic acid biosynthesis protein. | Metabolism |
| *SAUPAN002689000* | Backbone | 11 | Putative RNase adaptor protein RapZ. | Phage resistance |
| *frp* | Backbone | 12 | NAD(P)H-flavin oxidoreductase. Cell-wall protein linked to iron-restricted growth conditions. | Metabolism |
| *grlA* | Backbone | 14 | DNA topoisomerase IV subunit A. Fluoroquinolone resistance. | AMR |
| *mprF* | Backbone | 10 | Phosphatidylglycerol lysyltransferase, linked to methicillin resistance. | AMR |
| *pknB* | Backbone | 10 | Serine/threonine-protein kinase. | Metabolism |
| *SAOUHSC_01130* | Backbone | 13 | YfcC family arginine/ornithine APC transporter. | Metabolism |
| *pycA* | Backbone | 11 | Putative pyruvate carboxyl transferase. | Metabolism |
| *glyA* | Backbone | 11 | Serine hydroxymethyltransferase. | Metabolism |
| *fdhA* | Backbone | 13 | Formate dehydrogenase. Contains FeS cluster. | Metabolism |
| *rpoB* | Backbone | 22 | DNA-directed RNA polymerase subunit β. Rifampicin resistance. | AMR |
| *ponA* | Backbone | 24 | PBP2, cell wall biosynthesis. B-lactam-resistance. | AMR |
| *sucA* | Backbone | 14 | 2-oxoglutarate dehydrogenase E1 component. | Metabolism |
| *dinG* | Backbone | 14 | ATP-dependent helicase. | Metabolism |
| *dnaK* | Backbone | 14 | Molecular chaperone. | Metabolism |
| *pbpB* | Backbone | 15 | Penicillin binding protein 2, related to β-lactam resistance. | AMR |
| *atl* | Backbone | 17 | Bifunctional autolysin. | Virulence |
| *hsdS1* | Backbone | 14 | Restriction endonuclease subunit S. | Metabolism |
| *sdrH* | Backbone | 27 | Hypothetical Serine-Aspartate Repeat protein. Highly variable. | Metabolism |
| *lpdA* | Backbone | 9 | Dihydrolipoamide dehydrogenase. | Metabolism |
| *ybbH* | Backbone | 9 | MurR/RpiR family transcriptional regulator. | Metabolism |

**Supplementary Table 7.** Candidate genes in ST239 that show potential evidence of parallel evolution. Gene names and functions were identified in AureoWiki[v].

| ST | Isolate ID | β-lactam (ampicillin/penicillin) | β-lactam (oxacillin) | Carboxylic Acid (mupirocin) | Cephalosporin (ceftarolin) | Fosfomycin | Fusidic Acid | Glycopeptide (teicoplanin) | Glycopeptide (vancomycin) | Glycylcycline (tigecyclin) | Macrolide (erythromycin) | Oxazolidinone (linezolid) | Quinolone (moxifloxacin) | Rfamycin (rifampicin) | Streptogramin (synercid) | Diaminopyrimidine (trim/sulf) | Total number of resistance phenotypes |
|---|---|---|---|---|---|---|---|---|---|---|---|---|---|---|---|---|---|
| ST239 | ST239A | R | R | R | S | R | S | S | S | S | R | S | R | R | S | R | 9 |
| ST239 | ST239B | R | R | S | S | R | S | S | S | S | R | S | R | R | S | R | 8 |
| ST239 | ST239C | R | R | S | S | S | S | S | S | S | R | S | R | S | S | R | 6 |
| ST239 | ST239D | R | R | S | S | S | S | S | S | S | R | S | R | S | S | R | 6 |
| ST8 | ST8A | S | S | S | S | S | S | S | S | S | S | S | S | S | S | S | 0 |
| ST8 | ST8B | S | S | S | S | S | S | S | S | S | S | S | S | S | S | S | 0 |
| ST8 | ST8C | R | R | S | S | S | S | S | S | S | S | S | R | S | S | S | 3 |
| ST8 | ST8D | R | R | S | S | S | S | S | S | S | R | S | R | S | S | R | 5 |
| ST8 | ST8E | R | R | S | S | S | S | R | R | S | R | S | R | S | S | R | 8 |
| ST8 | ST8F | R | S | S | S | S | R | S | S | S | S | S | S | S | S | S | 2 |
| ST30 | ST30A | R | R | S | S | S | S | S | S | S | S | S | S | S | S | S | 2 |
| ST30 | ST30B | R | S | S | S | S | S | S | S | S | S | S | S | S | S | S | 1 |
| ST30 | ST30C | R | R | S | S | S | S | S | S | S | R | S | R | S | S | S | 4 |
| ST30 | ST30D | R | R | S | S | S | S | S | S | S | R | S | R | S | S | S | 4 |
| ST30 | ST30E | R | R | S | S | S | S | S | S | S | R | S | R | S | S | R | 6 |

**Supplementary Table 8.** AMR phenotypes for each isolate; susceptible, S; resistant, R.

i Sprouffske K, Wagner A. Growthcurver: an R package for obtaining interpretable metrics from microbial growth curves. BMC Bioinformatics. 2016;17. doi:10.1186/s12859-016-1016-7

ii Wellcome Sanger Institute. Major lineages of healthcare-associated methicillin-resistant Staphylococcus aureus in Singapore. 2011. Available from: https://www.ebi.ac.uk/ena/data/view/PRJEB2295

iii Robinson DA, Enright MC. Evolutionary Models of the Emergence of Methicillin-Resistant Staphylococcus aureus. AAC. 2003;47: 3926–3934. doi:10.1128/aac.47.12.3926-3934.2003

iv Wellcome Sanger Institute. Evolution of MRSA in a single institution over a 12 year period. 2012. Available from: https://www.sanger.ac.uk/resources/downloads/bacteria/staphylococcus-aureus.html#project_1851

v Fuchs S, Mehlan H, Bernhardt J, Hennig A, Michalik S, Surmann K, et al. Aureo Wiki‑The repository of the Staphylococcus aureus research and annotation community. International Journal of Medical Microbiology. 2018;308: 558–568. doi:10.1016/j.ijmm.2017.11.011
